# Inclusivity in fNIRS Studies: Quantifying the Impact of Hair and Skin Characteristics on Signal Quality with Practical Recommendations for Improvement

**DOI:** 10.1101/2024.10.28.620644

**Authors:** Meryem A. Yücel, Jessica E. Anderson, De’Ja Rogers, Parisa Hajirahimi, Parya Farzam, Yuanyuan Gao, Rini I. Kaplan, Emily J. Braun, Nishaat Mukadam, Sudan Duwadi, Laura Carlton, David Beeler, Lindsay K. Butler, Erin Carpenter, Jaimie Girnis, John Wilson, Vaibhav Tripathi, Yiwen Zhang, Bettina Sorger, Alexander von Lühmann, David C. Somers, Alice Cronin-Golomb, Swathi Kiran, Terry D. Ellis, David A. Boas

## Abstract

Functional Near-Infrared Spectroscopy (fNIRS) holds transformative potential for research and clinical applications in neuroscience due to its non-invasive nature and adaptability to real-world settings. However, despite its promise, fNIRS signal quality is sensitive to individual differences in biophysical factors such as hair and skin characteristics, which can significantly impact the absorption and scattering of near-infrared light. If not properly addressed, these factors risk biasing fNIRS research by disproportionately affecting signal quality across diverse populations. Our results quantify the impact of various hair properties, skin pigmentation as well as head size, sex and age on signal quality, providing quantitative guidance for future hardware advances and methodological standards to help overcome these critical barriers to inclusivity in fNIRS studies. We provide actionable guidelines for fNIRS researchers, including a suggested metadata table and recommendations for cap and optode configurations, hair management techniques, and strategies to optimize data collection across varied participants. This research paves the way for the development of more inclusive fNIRS technologies, fostering broader applicability and improved interpretability of neuroimaging data in diverse populations.

## 1 Introduction

Wearable technologies are rapidly becoming part of everyday life, revolutionizing how we monitor health and behavior ^1,2^. From fitness trackers to sleep monitors, these devices offer continuous, real-time insights into various physiological parameters. With ongoing advancements in hardware, Functional Near-Infrared Spectroscopy (fNIRS), a non-invasive functional brain imaging technology that measures cerebral hemodynamic responses, is poised to become a practical tool for brain monitoring in daily life, with significant implications for brain-computer interfaces, personalized medicine, and neurorehabilitation and the study of cognitive, developmental, and social neuroscience, and sports medicine among others ^3–10^. As fNIRS moves beyond laboratory settings into real-world applications, it is essential that the technology is accessible for all individuals. Ensuring inclusivity will maximize fNIRS’ impact globally, avoiding biases in data collection and making this technology beneficial to diverse populations ^11,12^.

The optical nature of fNIRS measurements makes the signal quality particularly susceptible to biophysical factors, especially hair and skin properties ^13^, as they can influence the absorption and scattering of near-infrared light ^11^. For example, more light is absorbed by dense hair, reducing the amount of near-infrared light that penetrates through the scalp and skull and reaches into the brain, thereby limiting the amount of light reflected back to the detectors. Similarly, more light is absorbed by darker hair as previously reported by Huang and colleagues ^14^. The darker skin i.e. skin with higher skin pigmentation also absorbs more near-infrared light ^13,15^. Moreover, the coupling of optodes to the scalp can be interfered with by hair characteristics such as density and type (i.e. the natural hair characteristics such as straight, wavy, curly and kinky) and thereby result in decreased signal quality. This poses a critical challenge to recruitment and data collection within fNIRS studies, as it potentially limits its use across diverse populations. To improve inclusivity in fNIRS measurements, it is crucial to tackle the considerations related to diverse hair and skin characteristics ^16^. Advancing optode design and implementing standardized procedures to account for natural variations in hair and skin characteristics are essential for optimizing fNIRS signal quality and overcoming related challenges. These efforts will make fNIRS technology more applicable and beneficial to a broader range of individuals and research settings, ensuring that it can be used effectively across diverse populations.

In this study, 115 participants were recruited to investigate the relationship between the presence of particular participant-level factors (i.e., hair and skin characteristics, head size, sex, and age) and fNIRS signal quality. Measurements were conducted using a continuous-wave fNIRS system with an optode array partially covering, fronto-temporo-parietal, and occipital cortices. Detailed protocols for skin and hair assessments were employed, including skin pigmentation (Melanin Index) measured by a melanometer ^17^ and high-resolution trichoscopy imaging. Participants completed resting state and motor tasks, and signal quality was optimized through cap adjustments. Correlation and regression analyses were performed to examine the relation between hair and skin characteristics, head size, sex and age and fNIRS signal quality, controlling for multiple comparisons and multicollinearity. In our discussion, we offer recommendations for best practices and guidelines to optimize fNIRS data collection across a diverse range of participant-level factors, driving forward a more inclusive approach to fNIRS research and application. In addition, we provide a suggested metadata table for researchers to include in their shared datasets to facilitate the expansion of our initial dataset. This will enable future studies to quantify the impact of diverse participant-level factors on fNIRS signal quality across larger populations through meta-analyses of multiple shared datasets. Such metadata will ensure that the influence of variables like hair and skin characteristics can be systematically evaluated, promoting a better understanding of how these factors affect fNIRS measurements, ultimately improving the generalizability of research findings.

## 2 Methods

### 2.1 Study Population

We recruited 115 participants (63 female, 51 male, 1 not reported; mean age: 26.4 years, SD: 10.3) under protocol #4502, which was reviewed and approved by the Boston University Institutional Review Board. All participants provided signed informed consent. The inclusion criteria required participants to be aged 18-89 years, have no self-reported history of neurological trauma or neurological or psychiatric disorders, not be taking any psychoactive medications, and be able to give written informed consent. The study sample included individuals who identified as of Asian descent (n = 45), European or Caucasian descent (n = 37), African descent (n = 27), those identifying as more than one race (n = 3), individuals not reporting their race (n = 2), and those categorized as other (n = 1).

### 2.2 fNIRS Measurements

fNIRS measurements were collected using a 3D-printed NinjaCap ^18^, a hexagonally netted cap made from NinjaFlex material (NinjaTek, Lititz PA, USA), populated with an optode array that covered the following three regions (Figure 1):

**Figure 1.**
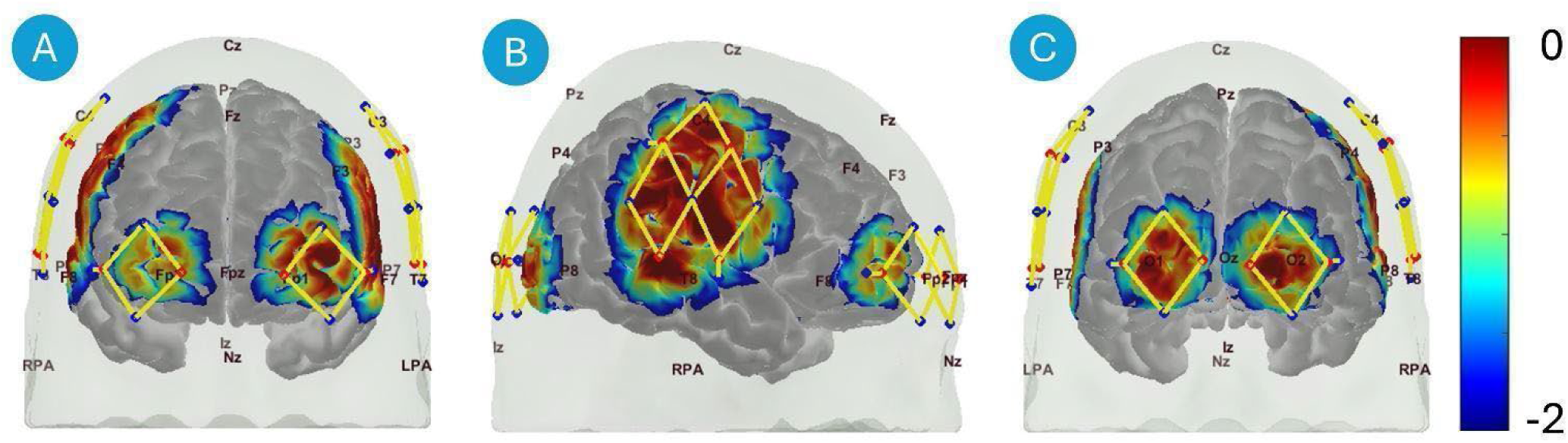
Optode array design. Anterior view (A), right view (B) and posterior view (C). Optode array design with sources (red dots), detectors (blue dots), and channels (yellow lines). Standard 10-20 locations are marked on the head for reference. Probe sensitivity is visualized on the brain surface, with values represented in log10 units.

**Forehead:** Dorsolateral prefrontal cortex is covered by four long-separation channels and one short-separation channel bilaterally at each hemisphere.
**Sides of the Head (Side):** Fronto-temporo-parietal cortex is covered by ten long-separation channels and two short-separation channels bilaterally at each hemisphere.
**Back of the Head (Back):** Occipital cortex is covered by four long-separation channels and one short-separation channel bilaterally at each hemisphere. The rationale for covering these regions was twofold: (1) to sample areas with varying hair distributions, such as the hairless forehead, and (2) to target functional brain regions involved in motor tasks, enabling us to assess how participant-level factors influence evoked brain responses. Long-separation channels were approximately 30 mm and short-separation channels were approximately 8 mm. The data were acquired using two continuous-wave fNIRS systems (NIRSport2, NIRx Medical Technologies, Germany), enabling the use of 16 dual-tip sources, 16 dual-tip detectors, and 4 short-separation detectors. Measurements were taken at two wavelengths (760 and 850 nm) with a sampling rate of 10.2 Hz. Please note that each hemisphere had a different type of detector, and only data from the right hemisphere, which used SiPD detectors, was analyzed throughout the paper. The left hemisphere has APD detectors from NIRx, and that data is available in our shared dataset, but we leave that for others to analyze in the future.

#### 2.2.1 Cap Placement

The cap placement procedure involved the following steps: The forehead was cleaned using alcohol pads. Velcro attachments were used on the top of the chair to secure the splitter boxes of the optode bundles, stabilizing the primary weight of the bundle cables. A metallic, adjustable cable management arm held the wires to prevent strain and reduce pressure on the participant’s head. The cap, sized either 55 cm or 57 cm depending on the participant’s head circumference, was placed on the participant’s head starting with the front part of the cap, and gently extending toward the back of the head. This front-to-back placement directionality prevents hair from falling forward under the optodes. To ensure consistent cap positioning across participants, the Cz marker on the cap was located midway between the nasion and inion, and equidistant from ear-to-ear. A chin strap was securely attached to stabilize the cap.

Before the first resting run, a brief initial optode-scalp coupling optimization process (<1 min) was performed which involved minor adjustments or ‘wiggling’ of optodes to establish preliminary optode-scalp coupling (fast capping) prior to running the acquisition software Aurora’s Signal Optimization function. This step was rudimentary and intentionally not the standard optimization procedure. Prior to the second resting run, the experimenters were instructed to make thorough adjustments of the cap and hair to improve signal quality (proper capping), which was guided by the continuous monitoring feature in Aurora. Scalp-coupling enhancing techniques while the cap is on include using cotton-tipped applicators to push hair from under optodes to the side. As needed, small amounts of hair or ultrasound gel were also applied via applicator directly under the optode; the NIRx optode may be temporarily removed from the grommet, gel applied to the grommet center as hair is pushed aside circularly, and optode replaced into the grommet. The rationale behind collecting resting-state data both after fast capping and proper capping was to enable a direct comparison of signal improvements resulting from thorough capping adjustments.

After achieving optimal signal quality, environmental adjustments were made to minimize interference from ambient light on intensity measurements. These adjustments involved turning off overhead lights (pulse-wave modulated LEDs), turning on floor lamps (incandescent bulbs), and placing an opaque shower cap over the fNIRS cap to block additional light sources, such as light from the computer monitor. Before starting the fNIRS recordings, the Signal Optimization function was run to verify that optimal signal quality was maintained.

### 2.3 Experimental Paradigms

Participants completed the following tasks: two runs of three-minute resting state and one run of a six-minute ball-squeezing task.

The first resting-state data was collected after fast capping, and the second resting-state data was collected after proper capping. During the resting-state measurement, participants were instructed to remain as still and mentally idle as possible.

Please note that for the signal quality assessments, we used the resting-state data after proper capping (second resting-state data). To see the signal improvement with capping, we used the resting-state data after fast capping and the resting data after proper capping.

For the six-minute left- and right-hand ball-squeezing task ^19^, participants sat in front of a computer screen, holding a rubber ball in each hand. At the start of the task, participants squeezed a ball in either their left or right hand at a self-selected frequency in response to visual cues presented on the screen, which included written words and an accompanying audio signal. They continued the squeezing motion until instructed to stop. Each stimulus lasted five seconds, with rest intervals randomly varying between 5 and 15 s. Each run consisted of 15 trials for both the left-hand and right-hand ball-squeezing conditions, presented in a fully randomized order. The stimulus presentation was created using PsychoPy software^20^.

### 2.4 Skin Measurements

#### Skin Pigmentation

A Skintone Pen (TPS 20, Courage-Khazaka, Cologne, Germany) was utilized for quantifying the skin pigmentation as Melanin Index. The Melanin Index is measured based on reflectance spectrophotometry at two wavelengths (660 and 880 nm) assuming that the main absorbers in the skin are melanin and hemoglobin, and is scaled between 0-99, with larger numbers indicative of darker skin pigmentation ^21,22^. The inner forearm, a region with minimal hair, was selected for measurements due to its comparable melanin levels to the scalp, which typically experiences less sun exposure. The area was cleaned using an alcohol wipe to ensure optimal contact. Placing the probe surface flush against the skin, pressure was applied until a distinct beep was sent by the device indicating that sufficient pressure had been applied. Measurements were repeated three times to obtain a more reliable average value.

#### Skin Type

Skin type was assessed using the Fitzpatrick Skin Phototype Scale, which classifies skin types based on sensitivity to ultraviolet (UV) radiation and tanning response ^23^. Skin type was visually categorized by the experimenter and noted based on a predefined regrouping of the Fitzpatrick Scale into three main categories: (1) Type I: Very fair skin and Type II: Fair skin, (2) Type II: Medium skin and light brown skin and (3) Type V: Brown skin, Type VI: Dark brown or black skin.

Please note that we used skin pigmentation in our regression analysis as it is a more objective way of quantifying skin pigmentation.

### 2.5 Hair Measurements

Multiple hair characteristics were measured using a hand-held dermatoscope (Handyscope) which is commonly used in dermatology research ^24^ (Handyscope; Dermlite, California, USA, and FotoFinder, Bad Birnbach, Germany) and subsequently assessed by TrichoLAB (Warsaw, Poland). The Handyscope was utilized to capture high-resolution photos of the scalp via the TrichoLab App on a smartphone, following the instructions provided by TrichoLab. The polarized light option of the system was employed. A comb was used to create a precise parting line on the scalp. The lens was fully attached to the scalp to prevent any light leakage from interfering with the trichoscopy images. Multiple high-resolution images from the scalp on the side of the head and on the back of the head were obtained. These images were securely transferred to the TrichoLAB server for evaluation by a TrichoLAB physician. The resultant report consisted of the following metrics provided for each region:

#### Average Number of Hairs (N/cm²)

Average number of hairs is the number of hairs normalized to one square centimeter.

#### Average Hair Shaft Thickness (µm)

Hair shafts visualized on 50X magnified images are subject to a diameter measurement using proprietary TrichoLAB algorithms with a precision of one micron. Hair dyeing does not influence the result. Average hair shaft thickness is the mean across the whole field of view.

#### Thin, Middle, and Thick Hair (%)

Categorized based on hair shaft thickness, this metric delineates the percentage of hairs falling into thin (≤30 µm), middle (30–50 µm), and thick (≥50 µm) categories within the observed field of view.

#### Single, Double, and Triple Follicular Units (%)

Utilizing follicular unit aggregation, this analysis discerns the ratio between single and multi-hair units. The outcome reveals the percentage distribution of single, double, and triple follicular units.

#### Cumulative Hair Thickness (mm/cm²)

Representing the sum of all hair diameters within one square centimeter, this metric amalgamates hair density and diameter, offering insights into scalp coverage and overall hair thickness/volume. This metric is calculated as the multiplication of the average number of hairs [N/cm2] and the average hair shaft thickness [μm] and 10^-3^ (to convert μm into mm).

#### Derived Sinclair Scale

This evaluation involves a rapid, instrument-free assessment of the width of the central hair parting line using visual cues, contributing to the qualitative analysis of hair health and progression according to the Sinclair Scale ^25^.

Additionally, we visually classified hair color, type and texture. The standards used include the Fischer-Saller Scale, Andre Walker Hair Typing System, and the FIA Hair Typing System (Table 1).

**Table 1.**
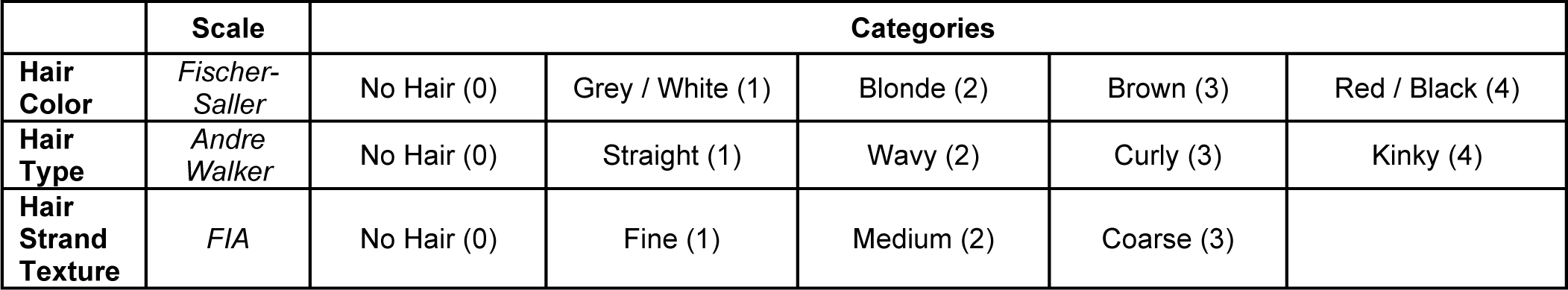
Classification scales for hair characteristics including hair color, type, and strand texture. Hair color is categorized using the Fischer-Saller scale, hair type follows the Andre Walker classification, and hair strand texture is assessed using the FIA scale.

#### Hair Color

The Fischer-Saller Scale categorizes hair color into four main types. Category 1 includes individuals with grey or white hair ^26^. Category 2 encompasses those with blonde hair. Category 3 includes individuals with brown hair. Category 4 covers individuals with red or black hair. Please note that the rationale for combining the red and black hair into one category is that the pheomelanin, which is a subgroup of melanin pigments that gives red hair its color, absorbs more light in the near-infrared spectrum than eumelanin (the pigment in the black hair) ^27,28^. In the case of no hair, the value was set to 0.

#### Hair Type

Hair type refers to the natural structure of hair e.g. curl pattern. The Andre Walker Hair Typing System classifies hair based on its curl pattern. This system identifies four distinct types of hair: Type 1 (straight), Type 2 (wavy), Type 3 (curly), and Type 4 (kinky) ^29^. In the case of no hair, the value was set to 0.

#### Hair Strand Texture

Hair texture refers to the thickness or coarseness of individual hair strands. The FIA Hair Typing System categorizes hair based on its thickness or coarseness. Type 1 hair is classified as fine, Type 2 as medium, and Type 3 as coarse. In the case of no hair, the value was set to 0.

### 2.6 Head Measurements

Measurements are taken around the head, including head circumference, nasion to inion distance, and ear to ear distance.

#### Head Circumference

Head circumference represents the measurement taken around the widest part of the head, encircling the cranium from the forehead, around the back, and returning to the starting point.

#### Nasion to Inion

Nasion to inion is a measurement taken from the nasion, the midpoint between the eyes where the frontal bone and nasal bones meet, to the inion, which is the prominent bony protuberance at the base of the occipital bone on the skull’s posterior. This measurement involves crossing Cz, which is a specific scalp location according to the international 10-20 system of EEG electrode placement.

#### Ear to Ear

This measurement is taken horizontally between the left and right preauricular points, which are the bony protrusions located just in front of each ear. The distance also crosses the vertex of the head (Cz).

### 2.7 Combined Hair-Skin Metric

#### Combined Hair-Skin Metric

The combined hair-skin metric was calculated using data from the Side and the Back of the Head to capture the joint impact of hair and skin properties. Four representative metrics were selected: average hair-shaft thickness (range: 42–88 µm), hair type (scale: 0–4), hair color (scale: 0–4), and skin pigmentation (range: 4.7–90). These measures were chosen to reflect key structural and pigmentation characteristics relevant to fNIRS signal quality. The values of each of these metrics were initially normalized to a range of [0, 1] using the formula below where min and max are the minimum and maximum of the range listed in parentheses above:

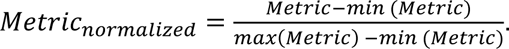

Subsequently, for each subject, the average of the normalized values of these four metrics, indexed with *i* below, where *m* = 4, is calculated as to produce a single combined metric as below:

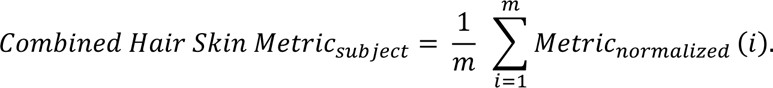

### 2.8 Participant-Level Factors

The mean and standard deviation of measured hair characteristics across participants at the Side and the Back of the Head are displayed in Table 2 and skin and hair characteristics for the whole head are displayed in Table 3.

**Table 2:**
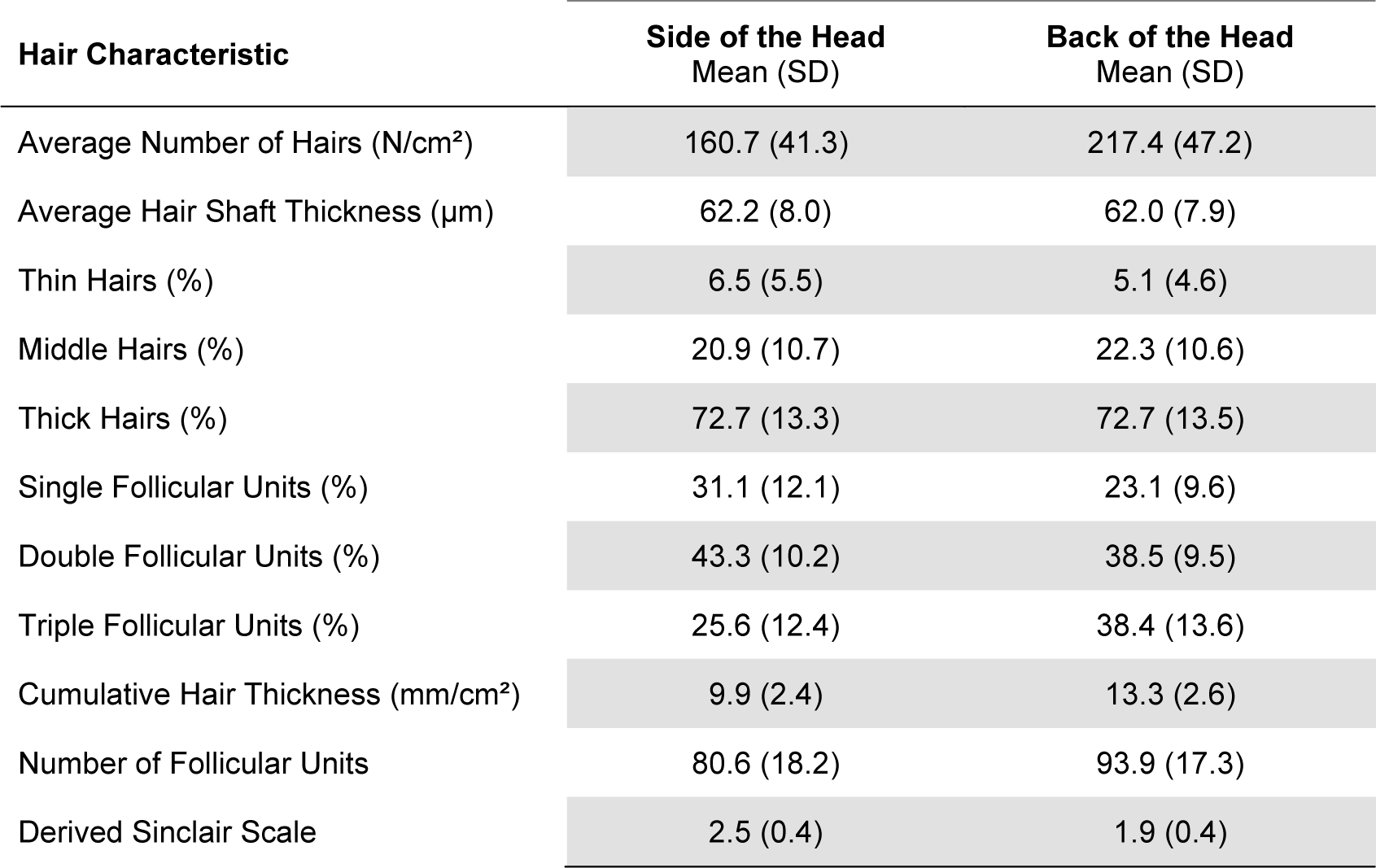
Mean and standard deviation of hair characteristics on the Side and Back of the Head.

| Hair Characteristic | Side of the Head<br>Mean (SD) | Back of the Head<br>Mean (SD) |
| --- | --- | --- |
| Average Number of Hairs (N/cm<sup>2</sup>) | 160.7 (41.3) | 217.4 (47.2) |
| Average Hair Shaft Thickness (μm) | 62.2 (8.0) | 62.0 (7.9) |
| Thin Hairs (%) | 6.5 (5.5) | 5.1 (4.6) |
| Middle Hairs (%) | 20.9 (10.7) | 22.3 (10.6) |
| Thick Hairs (%) | 72.7 (13.3) | 72.7 (13.5) |
| Single Follicular Units (%) | 31.1 (12.1) | 23.1 (9.6) |
| Double Follicular Units (%) | 43.3 (10.2) | 38.5 (9.5) |
| Triple Follicular Units (%) | 25.6 (12.4) | 38.4 (13.6) |
| Cumulative Hair Thickness (mm/cm<sup>2</sup>) | 9.9 (2.4) | 13.3 (2.6) |
| Number of Follicular Units | 80.6 (18.2) | 93.9 (17.3) |
| Derived Sinclair Scale | 2.5 (0.4) | 1.9 (0.4) |

**Table 3:** Mean and standard deviation of whole-head hair characteristics, skin properties, and head measurements for the study population.

| Participant-Level Factors | Mean (SD) |
| --- | --- |
| Skin Pigmentation | 25.3 (18.1) |
| Skin Type | 2.0 (0.9) |
| Hair Color | 3.2 (1.0) |
| Hair Type | 1.8 (1.1) |
| Hair Strand Texture | 1.7 (0.8) |
| Head Circumference (cm) | 57.1 (2.3) |
| Nasion to Inion (cm) | 35.4 (2.4) |
| Ear to Ear (cm) | 35.8 (2.7) |

### 2.9 Signal Quality Metrics

We used three metrics in this study: Uncorrected Signal Mean, Corrected Signal Mean and the Scalp Coupling Index (SCI).

#### Uncorrected Signal Mean

The mean of the raw signal intensity time course, averaged across two wavelengths. The Uncorrected Signal Mean is used in the Noise Equivalent Power (NEP) figure and for the comparison between the fast capping and proper capping.

#### Corrected Signal Mean

The mean of the raw signal intensity time course, averaged across two wavelengths, was divided by the LED power percentage of the source power (for each channel) from the NIRx configuration file, multiplied by 100, and then logarithmically transformed (log10). Source power correction was applied to facilitate comparison of signals across different channels and subjects by correcting for differences in source illumination power. This also enhanced the visibility of hair and skin effects, as without this correction, the values obtained (i.e. Uncorrected Signal Mean) are after the system has performed source modulation to reduce these effects. The logarithmic transformation approximately normalizes the distribution and helps approximate a linear relationship with independent variables. The Corrected Signal Mean is used in signal quality analysis (i.e. correlation analysis and multiple linear regression).

**Scalp Coupling Index (SCI)** is a metric used to evaluate the quality of signal acquisition and the contact between the optodes and the scalp surface ^30^. Higher SCI often indicates better optode-scalp coupling and superior signal quality. To compute SCI, the pulsatile cardiac waveform is extracted by band-pass filtering the fNIRS signal at both wavelengths between 0.5 and 2.5 Hz. The filtered signals are normalized to their standard deviation. Finally, SCI is computed as the zero-lag cross-correlation between the normalized signals at the two wavelengths. This calculation estimates the similarity between the two normalized signals and provides an index reflecting the quality of signal transmission and detection between the fNIRS optodes and the scalp surface. A Fisher z-transformation was applied to the SCI values in order to stabilize the variance and normalize the distribution of correlation coefficients. The SCI is used in signal quality analysis (i.e. correlation analysis and multiple linear regression).

### 2.10 Violin Plots

Violin plots are used to visualize both continuous independent variables (e.g., hair thickness) and categorical variables (e.g., hair color). Continuous variables were divided into four equal-width bins between the minimum and maximum values of the independent variable, facilitating a clearer visualization of the distribution across different ranges. However, in the correlation analysis as well as multiple linear regression analysis, these continuous variables were treated as continuous, not binned.

### 2.11 Correlation Analysis and Robust Linear Fitting

Correlation analyses were performed to investigate the relationships between participant-level factors and Corrected Signal Mean on the Forehead, Side of the Head and Back of the Head. The Spearman’s rank correlation coefficient (ρ) and the corresponding p-values were calculated for each participant characteristic using MATLAB’s *corr* function (R2023a, MathWorks Inc., Natick, MA). Multiple comparisons corrections were applied using the Benjamini-Hochberg procedure to control the false discovery rate ^31^. Correlations were conducted within specific ROIs, with Corrected Signal Mean correlated channel-wise with the corresponding hair characteristics measured in the same region. For instance, the Corrected Signal Mean in individual channels on the Side of the Head was correlated with hair characteristics measured at that location.

Robust linear regression was conducted to determine the relationship between each participant-level factor and the Corrected Signal Mean. The robust fitting method was chosen to minimize the influence of outliers on the regression results.

The fold change in the Corrected Signal Mean between the 1^st^ (lower bound, *L*) and 99^th^ (upper bound, *U*) percentiles of each participant characteristic was computed using the endpoints of the robust linear regression model and then transformed back to their original scale (unlogged, using the inverse of log10). If the upper bound exceeds or equals the lower bound, the fold change is calculated as the ratio of the upper bound to the lower bound. Conversely, if the lower bound exceeds the upper bound, the calculation is inverted to accurately reflect the direction of change.

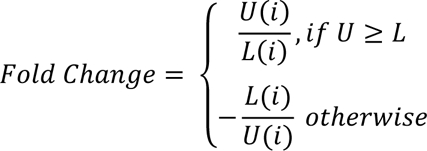

### 2.12 Multiple Linear Regression Analysis

Multiple linear regression analysis with robust regression was conducted to explore the relationship between participant-level factors and the signal-quality metrics during the second resting-state run (i.e. after proper capping) using the built-in function *fitlm* in MATLAB R2023a (MathWorks Inc., Natick, MA). The dependent variable (Y) was the signal quality metrics, and the independent variables were participant-level factors.

To evaluate multicollinearity among independent variables, Pearson correlation coefficients were calculated between pairs of independent variables using the *corr* function in MATLAB. The correlation matrix was examined to identify highly correlated variables with correlation coefficients exceeding a predefined threshold (0.5). If two independent variables had a correlation higher than 0.5, we only kept one of them in the regression model. Our decision of choice between the highly correlated independent variables was based on whether the metric is more objective and interpretable (e.g. skin pigmentation is chosen over skin type). We have calculated and reported the Variance Inflation Factor (VIF) which quantifies how much the variance of a regression coefficient is inflated due to multicollinearity with other predictors. High VIF (>10) indicates problematic multicollinearity. Moderate VIF (5-10) is acceptable in some contexts but generally should be monitored. Low VIF (<5) generally indicates low multicollinearity, which is desirable.

### 2.13 Statistical tests

We utilized two-sided Mann-Whitney U Tests to compare the fNIRS signal quality metrics between different ranges of participant-level factors. To account for multiple comparisons, the Benjamini-Hochberg correction was applied to the p-values obtained by the Mann-Whitney U Test and Correlation Analysis ^31^. We set the significance threshold after the correction to 0.01.

Multiple linear regression analyses were performed to identify significant predictors of fNIRS signal quality. The t-statistics for each predictor variable were computed to assess their individual contributions to the model. Additionally, effect sizes were calculated using Cohen’s f² to quantify the practical significance of each predictor ^32^. According to Cohen’s (1988) guidelines, f² ≥ 0.02, f² ≥ 0.15, and f² ≥ 0.35 represent small, medium, and large effect sizes, respectively.

To evaluate the improvement in signal quality following cap adjustments, paired t-tests were conducted. This analysis compared the signal quality measurements regarding the resting-state data before and after the second, thorough cap adjustments within the same participants, determining whether the adjustments resulted in statistically significant improvements.

### 2.14 General Linear Model

In order to analyze the data of the ball-squeezing experiment, the raw intensity measured at the motor cortex channels were converted into optical density (OD). The channels with SNR lower than 5 were excluded from further analysis. The data were then low-pass filtered with a sixth-order Butterworth filter at 0.5 Hz. The changes in OD signal were then converted into concentration changes of HbO_2_ and HbR by employing the modified Beer–Lambert law without pathlength correction. The hemodynamic response function (HRF) was estimated by a General Linear Model (GLM) that uses ordinary least squares estimation. The hemodynamic response function (HRF) was modeled using a modified gamma function with parameters tau, sigma, and duration set to [0.1, 3, 5] for HbO₂ and [1.8, 3, 5] for HbR, over a time range from −2 to 12 s. For each long-separation channel, the short-separation channel time course that had the highest correlation with that long-separation channel’s time course was used as a regressor in the GLM to model the systemic physiology in that long-separation channel ^33^. Baseline correction was performed by subtracting the mean of the HRF between −2 and 0 s from the rest of the HRF. The t statistics for the beta value for the HbO_2_ fit was obtained ^34^.

## 3 Results

### 3.1 Effects of Hair Characteristics on fNIRS Signal Quality

Figure 2 presents violin plots that demonstrate the relationship between hair characteristics and the Corrected Signal Mean on the Side of the Head. The data are divided into four equal-width bins based on the hair characteristics, with each bin depicted by a distinct color. The Mann-Whitney U Test after Benjamini-Hochberg correction reveals a significant increase in the Corrected Signal Mean on the Side of the Head from the lowest bin to the highest bin for Cumulative Hair Thickness (Figure 2C).

**Figure 2.**
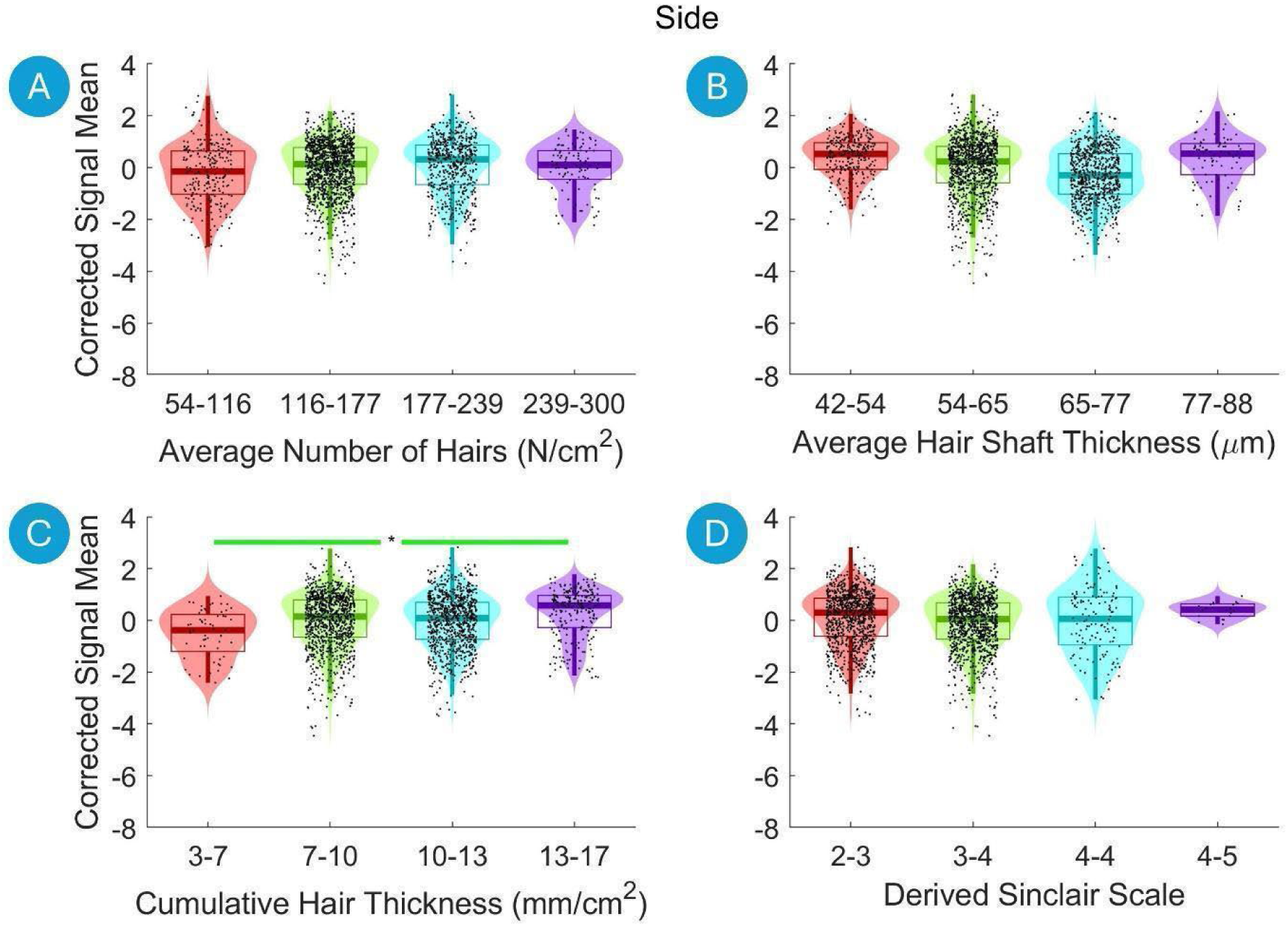
Violin plots illustrating the relationship between the hair characteristics and the Corrected Signal Mean on the Side of the Head. The data are divided into four equal-width bins based on the hair characteristics, with different colors representing each bin. Violin plots show the distribution of the fNIRS signal data for each bin, with boxplot elements overlayed to indicate quartiles, medians, and potential outliers. Significant differences between to first and the last bins are marked with asterisks (*, p < 0.01 after Benjamini-Hochberg correction). A green line indicates a statistically significant increase, and a red line indicates a statistically significant decrease.

Figure 3 displays violin plots that demonstrate the relationship between the hair characteristics and the Corrected Signal Mean on the Back of the Head. The Mann-Whitney U Test after Benjamini-Hochberg correction reveals a significant increase in the Corrected Signal Mean from the lowest hair density bin to the highest density bin when examining the Derived Sinclair Scale (Figure 3D). In contrast, there was a pronounced decrease in Corrected Signal Mean from the lowest to the highest bins for both Average Hair Shaft Thickness and Cumulative Hair Thickness (Figure 3B and C).

**Figure 3.**
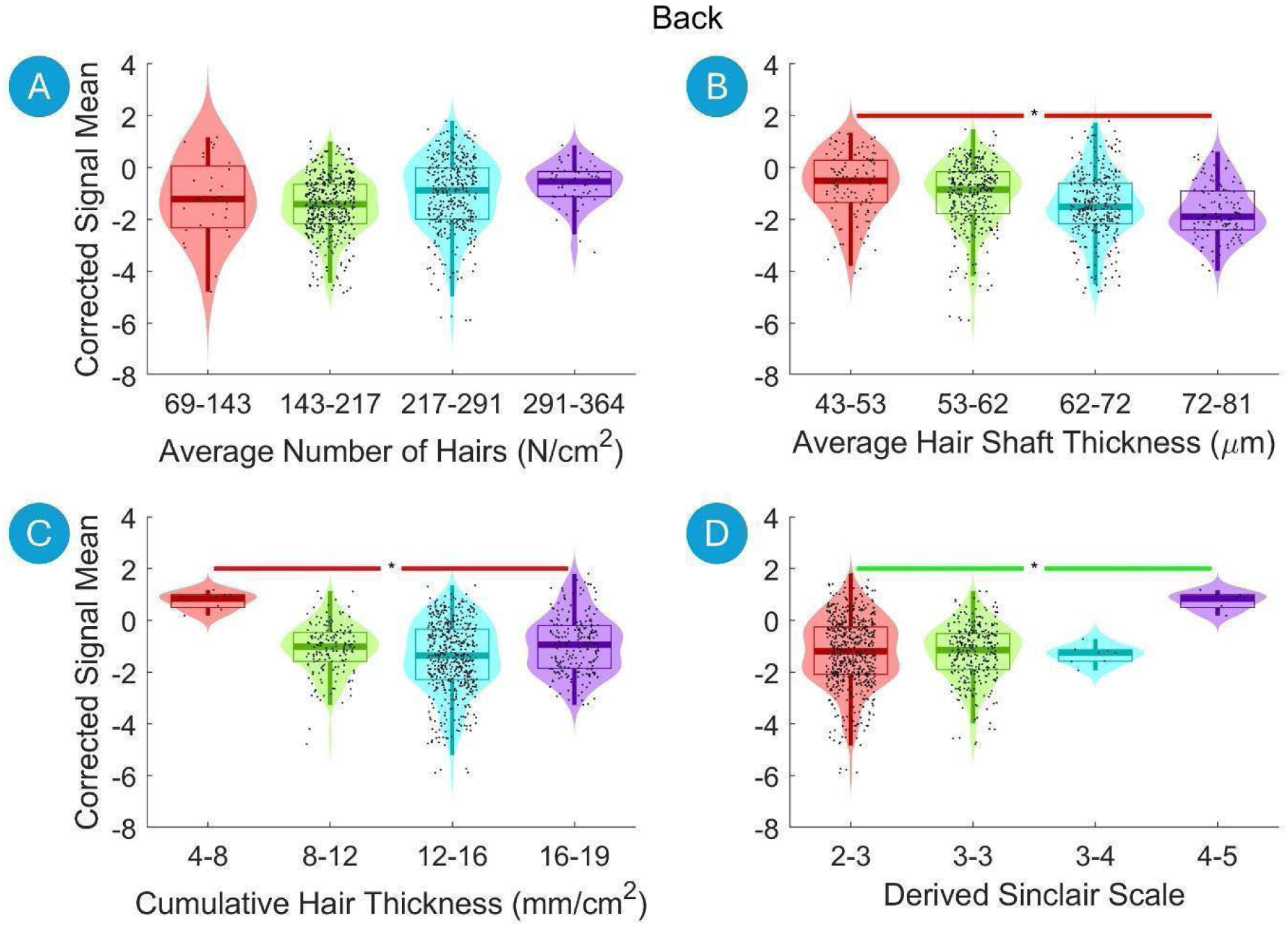
Violin plots illustrating the relationship between the hair characteristics and the Corrected Signal Mean on the Back of the Head. The data are divided into four equal-width bins based on the hair characteristics, with different colors representing each bin. Violin plots show the distribution of the fNIRS signal data for each bin, with boxplot elements overlayed to indicate quartiles, medians, and potential outliers. Significant differences between bins are marked with asterisks (*, p < 0.01 after Benjamini-Hochberg correction). A green line indicates a statistically significant increase, and a red line indicates a statistically significant decrease.

Figure 4 illustrates violin plots that depict the relationship between the categorical hair characteristics and the Corrected Signal Mean on the Side and Back of the Head, shown in the top and bottom rows, respectively. The analysis reveals a significant decrease in Corrected Signal Mean from individuals with no hair to those with black hair both on Side and Back of the Head (Figure 4A and D). Similarly, there is a significant reduction in Corrected Signal Mean from no hair to kinky hair, following a gradual decrease across Hair Type categories from straight, wavy, curly, to kinky (Figure 4B and E). Hair Strand Texture also exhibits a similar trend, where signal intensity significantly decreases from no hair to fine, medium, and coarse hair (Figure 4C and F).

**Figure 4.**
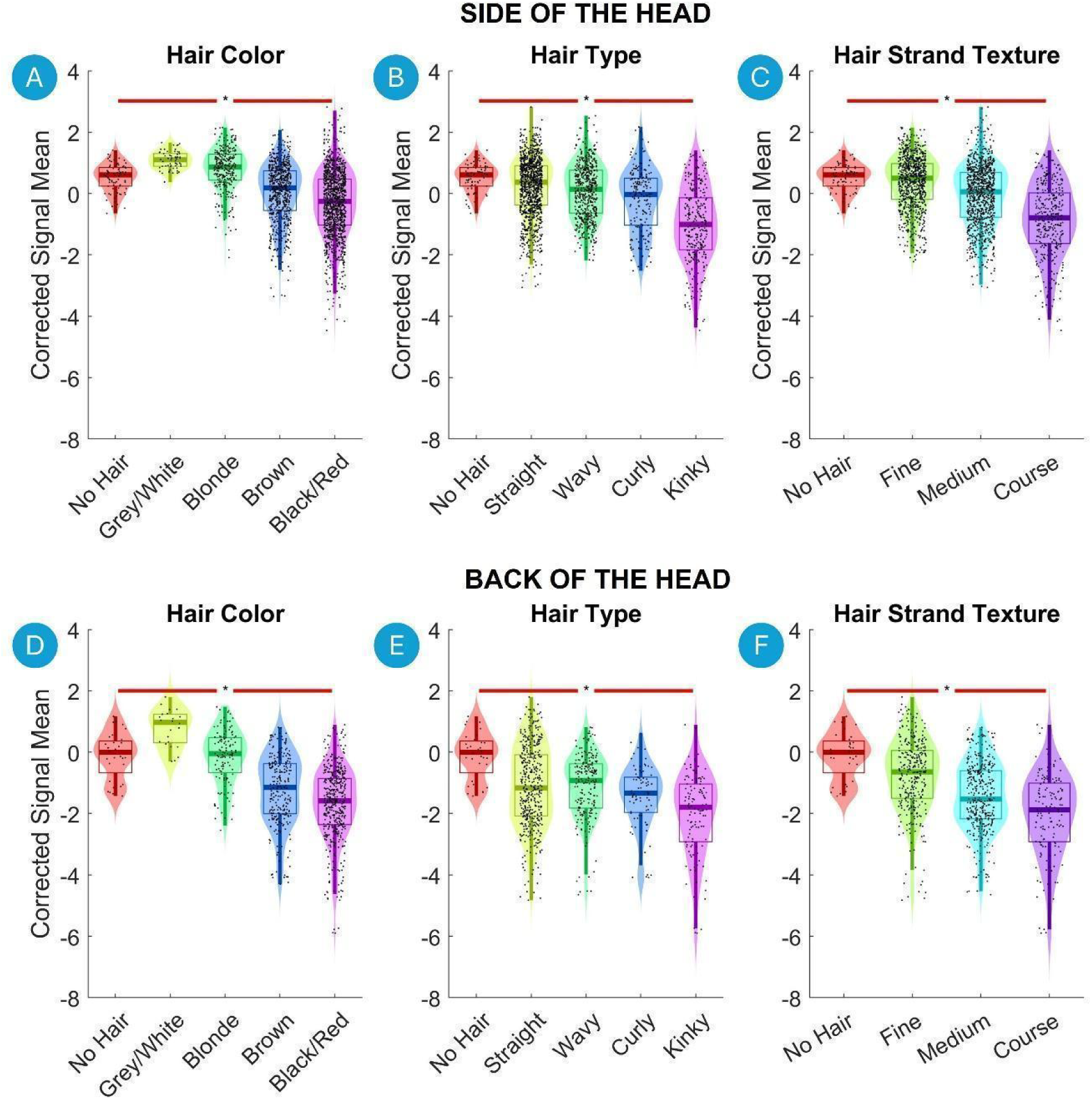
Violin plots illustrating the relationship between the hair characteristics and the Corrected Signal Mean on the Side and Back of the Head. Each category is represented with a different color. Violin plots show the distribution of the fNIRS signal data for each bin, with boxplot elements overlayed to indicate quartiles, medians, and potential outliers. Significant differences between the first and last bins are marked with asterisks (*, p < 0.01 after Benjamini-Hochberg correction). A green line indicates a statistically significant increase, and a red line indicates a statistically significant decrease.

### 3.2 Effects of Skin Characteristics on fNIRS Signal Quality

Figure 5 presents violin plots of the Corrected Signal Mean across three regions versus the Skin Pigmentation. The Corrected Signal Mean dropped significantly with increasing Skin Pigmentation on the Forehead, Side of the Head as well as on the Back of the Head (Figure 5A, B and C respectively).

**Figure 5.**
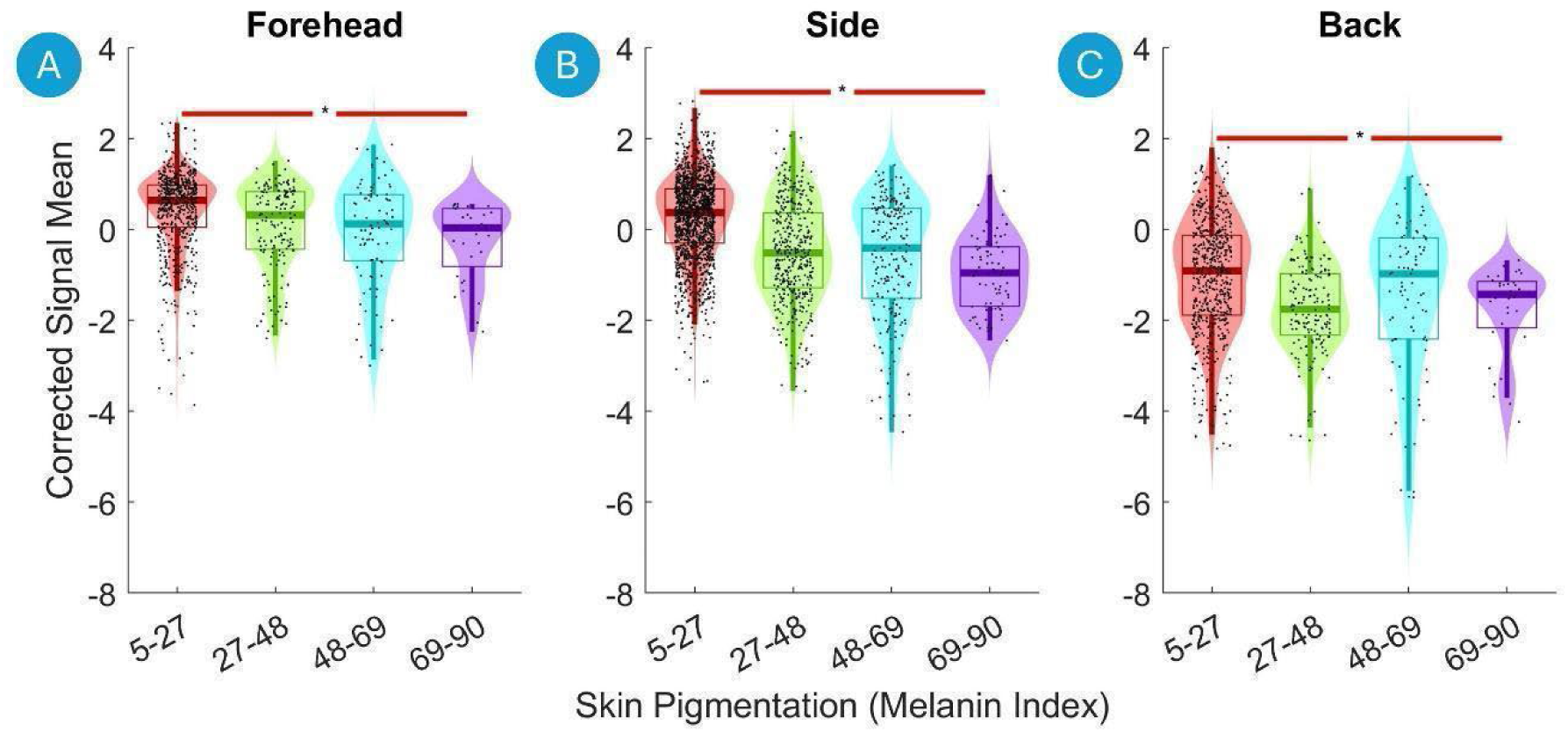
Violin plots of the Corrected Signal Mean on the Forehead, Side of the Head and Back of the Head versus the skin pigmentation. The data are divided into four equal-width bins based on the skin pigmentation, with different colors representing each bin. Violin plots show the distribution of the fNIRS signal data for each bin, with boxplot elements overlayed to indicate quartiles, medians, and potential outliers. Significant differences between the first and the last bins are marked with asterisks (*, p < 0.01 after Benjamini-Hochberg correction). A green line indicates a statistically significant increase, and a red line indicates a statistically significant decrease.

### 3.3 Effects of Head Size, Age and Sex on fNIRS Signal Quality

Figure 6 presents violin plots of the Corrected Signal Mean on the Forehead, the Side of the Head and the Back of the Head versus the head circumference (top row) and age (bottom row). The Corrected Signal Mean drops significantly with increasing head circumference on the Side the Head (Figure 6, top panels). Conversely, there is a significant increase in the detected Corrected Signal Mean with increasing age on the Forehead, Side of the Head and Back of the Head (Figure 6D, E and F respectively). Finally, there was a significant increase in Corrected Signal Mean from Male to Female on the Forehead and the Side of the Head (Figure 7A and B).

**Figure 6.**
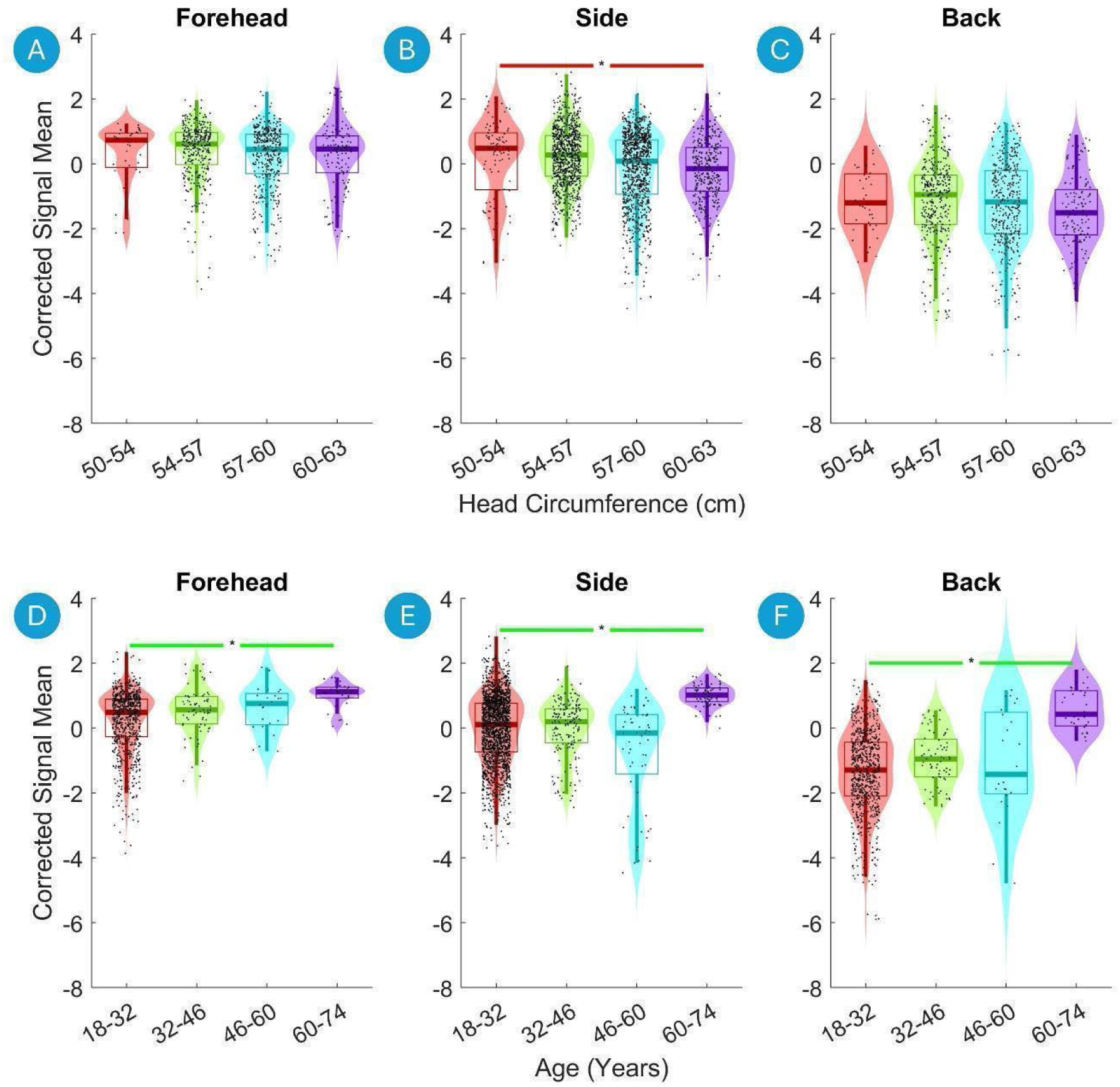
Violin plots of the Corrected Signal Mean on the Forehead, on the Side of the Head and on the Back of the Head versus head circumference (top) and age (bottom). The data are divided into four equal-width bins based on the head circumference and age, with different colors representing each bin. Violin plots show the distribution of the fNIRS signal data for each bin, with boxplot elements overlayed to indicate quartiles, medians, and potential outliers. Significant differences between the first and last bins are marked with asterisks (*, p < 0.01 after Benjamini-Hochberg correction). A green line indicates a statistically significant increase, and a red line indicates a statistically significant decrease.

**Figure 7.**
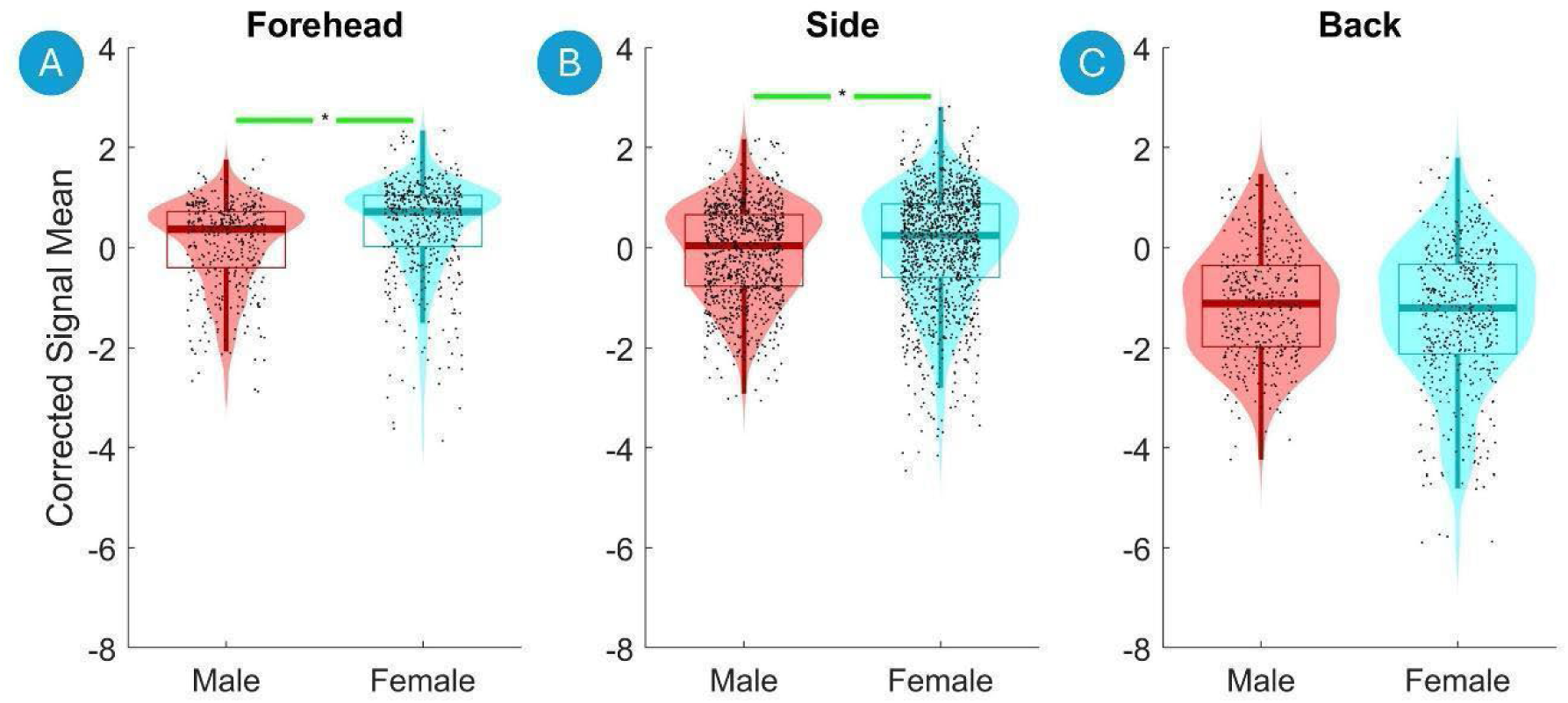
Violin plots of the Corrected Signal Mean on the Forehead, Side of the Head and Back of the Head versus sex. Violin plots show the distribution of the fNIRS signal data for each bin, with boxplot elements overlayed to indicate quartiles, medians, and potential outliers. Significant differences between the first and last bins are marked with asterisks (*, p < 0.01 after Benjamini-Hochberg correction), indicating statistical significance assessed using the Mann-Whitney U Test with Benjamini-Hochberg correction for multiple comparisons. A green line indicates a statistically significant increase, and a red line indicates a statistically significant decrease.

The correlation analysis, summarized in Table 4, reveals several significant relationships between various participant-level factors and the Corrected Signal Mean on the Forehead, Side of the Head and Back of the Head. These results are underscored by statistically significant correlation coefficients (ρ) with Benjamini-Hochberg corrected p-values, alongside substantial fold changes calculated between the 1st and 99th percentiles of the participant-level factors.

**Table 4:** Correlation Analysis of Corrected Signal Mean on the Forehead, Side of the Head and Back of the Head with participant-level factors. The fold change in signal quality measure calculated between the 1st and 99th percentiles of participant-level factors, the correlation coefficients (ρ), Benjamini-Hochberg corrected p-values for the correlation coefficients, and the slopes and intercepts of the linear fit are provided. Red highlights the statistically significant correlations of p<=0.01 after Benjamini-Hochberg correction.

| Forehead | Fold Change | rho | p-value | Slope | Intercept |
| --- | --- | --- | --- | --- | --- |
| Skin Pigmentation | -5 | -0.28 | 0.02 | -0.010 | 0.86 |
| Skin Type | -2 | -0.22 | 0.07 | -0.124 | 0.86 |
| Gender | 3 | 0.41 | <0.01 | 0.401 | 0.05 |
| Head Circumference | -2 | -0.12 | 0.30 | -0.031 | 2.38 |
| Nasion to Nasion | -3 | -0.23 | 0.07 | -0.044 | 2.17 |
| Ear to Ear | -3 | -0.18 | 0.12 | -0.043 | 2.15 |
| Age | 4 | 0.06 | 0.67 | 0.010 | 0.35 |

| Side of the Head | Fold Change | rho | p-value | Slope | Intercept |
| --- | --- | --- | --- | --- | --- |
| Average Number of Hairs | 4 | 0.16 | 0.18 | 0.003 | -0.07 |
| Average Hair Shaft Thickness | -2 | -0.15 | 0.20 | -0.009 | 0.93 |
| Thin Hairs Percent | -10 | -0.24 | 0.04 | -0.036 | 0.63 |
| Middle Hairs Percent | 4 | 0.09 | 0.41 | 0.010 | 0.16 |
| Thick Hairs Percent | -1 | 0.00 | 0.99 | -0.002 | 0.52 |
| Single Follicular Units Percent | -2 | -0.04 | 0.78 | -0.003 | 0.48 |
| Double Follicular Units Percent | 1 | 0.00 | 0.99 | 0.001 | 0.36 |
| Triple Follicular Units Percent | 2 | 0.07 | 0.54 | 0.004 | 0.29 |
| Cumulative Hair Thickness | 2 | 0.10 | 0.39 | 0.033 | 0.06 |
| Number of Follicular Units | 4 | 0.16 | 0.18 | 0.007 | -0.17 |
| Derived Sinclair Scale | -2 | -0.10 | 0.39 | -0.165 | 0.79 |
| Skin Pigmentation | -26 | -0.47 | <0.01 | -0.019 | 0.86 |
| Skin Type | -5 | -0.42 | <0.01 | -0.362 | 1.09 |
| Hair Color | -15 | -0.47 | <0.01 | -0.294 | 1.32 |
| Hair Type | -11 | -0.36 | <0.01 | -0.259 | 0.85 |
| Hair Strand Texture | -12 | -0.40 | <0.01 | -0.357 | 0.98 |
| Gender | 2 | 0.13 | 0.29 | 0.229 | 0.02 |
| Head Circumference | -3 | -0.15 | 0.19 | -0.045 | 2.94 |
| Nasion to Nasion | -3 | -0.18 | 0.14 | -0.046 | 2.00 |
| Ear to Ear | -8 | -0.23 | 0.04 | -0.071 | 2.91 |
| Age | 4 | 0.08 | 0.48 | 0.012 | 0.08 |

| Back of the Head | Fold Change | rho | p-value | Slope | Intercept |
| --- | --- | --- | --- | --- | --- |
| Average Number of Hairs | 13 | 0.25 | 0.02 | 0.005 | -1.93 |
| Average Hair Shaft Thickness | -22 | -0.30 | <0.01 | -0.040 | 1.54 |
| Thin Hairs Percent | -2 | -0.07 | 0.56 | -0.014 | -0.86 |
| Middle Hairs Percent | 32 | 0.30 | <0.01 | 0.032 | -1.64 |
| Thick Hairs Percent | -11 | -0.21 | 0.06 | -0.018 | 0.39 |
| Single Follicular Units Percent | 4 | 0.11 | 0.36 | 0.013 | -1.22 |
| Double Follicular Units Percent | -3 | -0.12 | 0.35 | -0.011 | -0.49 |
| Triple Follicular Units Percent | -1 | 0.02 | 0.83 | -0.001 | -0.90 |
| Cumulative Hair Thickness | 2 | 0.06 | 0.61 | 0.020 | -1.20 |
| Number of Follicular Units | 15 | 0.27 | 0.01 | 0.013 | -2.20 |
| Derived Sinclair Scale | -1 | -0.08 | 0.56 | -0.042 | -0.85 |
| Skin Pigmentation | -11 | -0.35 | <0.01 | -0.014 | -0.58 |
| Skin Type | -9 | -0.38 | <0.01 | -0.476 | 0.06 |
| Hair Color | -147 | -0.50 | <0.01 | -0.542 | 0.84 |
| Hair Type | -7 | -0.18 | 0.10 | -0.208 | -0.55 |
| Hair Strand Texture | -38 | -0.38 | <0.01 | -0.528 | -0.03 |
| Gender | 1 | 0.03 | 0.81 | 0.040 | -0.99 |
| Head Circumference | -2 | -0.07 | 0.56 | -0.019 | 0.14 |
| Nasion to Nasion | -1 | -0.07 | 0.56 | -0.007 | -0.68 |
| Ear to Ear | -4 | -0.14 | 0.28 | -0.049 | 0.84 |
| Age | 53 | 0.13 | 0.30 | 0.031 | -1.76 |

On the Forehead, Sex correlated positively, indicating that Female participants tend to have higher Corrected Signal Mean than Male participants. In addition, Skin Pigmentation showed close-to-significant (p = 0.02) negative correlation with Corrected Signal Mean, indicating that higher pigmentation levels are associated with Corrected Signal Mean.

On the Side of the Head, significant negative correlations were found with several hair characteristics. On the Side of the Head, Skin Pigmentation exhibited a strong negative correlation (ρ = −0.47, p < 0.01), with a substantial fold change of −26, indicating that higher pigmentation levels are associated with reduced Corrected Signal Mean. Hair Color also demonstrated a significant negative correlation (ρ = - 0.47, p < 0.01) and a notable fold change of −15, indicating that darker hair is linked to lower Corrected Signal Mean. Additionally, Hair Strand Texture presented a significant negative correlation (ρ = −0.40, p < 0.01) with a fold change of −12, suggesting that coarser hair types contribute to diminished Corrected Signal Mean. Skin Type reflected a significant negative correlation with Corrected Signal Mean (ρ = - 0.42, p < 0.01) and a fold change of −5. Lastly, Hair Type showed a significant negative correlation (ρ = - 0.36, p < 0.01) with Corrected Signal Mean.

On the Back of the Head, Hair Color exhibited a strong negative correlation (ρ = −0.50, p < 0.01) with a substantial fold change of −147, indicating that darker hair is linked to significantly lower Corrected Signal Mean. Hair Strand Texture presented a significant negative correlation (ρ = −0.38, p < 0.01) with a fold change of −38, implying that coarser hair types contribute to diminished Corrected Signal Mean. Average Hair Shaft Thickness also demonstrated a significant negative correlation (ρ = −0.30, p < 0.01) with a fold change of −22, suggesting that thicker hair may further reduce Corrected Signal Mean. Skin Pigmentation had a significant negative correlation (ρ = −0.35, p < 0.01) with a fold change of −11, indicating that higher pigmentation levels are associated with a decreased Corrected Signal Mean. Skin Type exhibited a significant negative correlation (ρ = −0.38, p < 0.01) with a fold change of −9. Conversely, the Middle Hairs Percent showed a significant positive correlation with Corrected Signal Mean (ρ = 0.30, p < 0.01) with a fold change of 32 and the Number of Follicular Units demonstrated a positive correlation (ρ = 0.27, p = 0.01) with a fold change of 15.

### 3.4 Multiple Regression Analysis of the Effect of Participant-level Factors on fNIRS Signal Quality

Multiple linear regression analysis was conducted to identify significant predictors of Corrected Signal Mean at the Forehead, Side of the Head and Back of the Head (Table 5, Figure 8).

**Figure 8.**
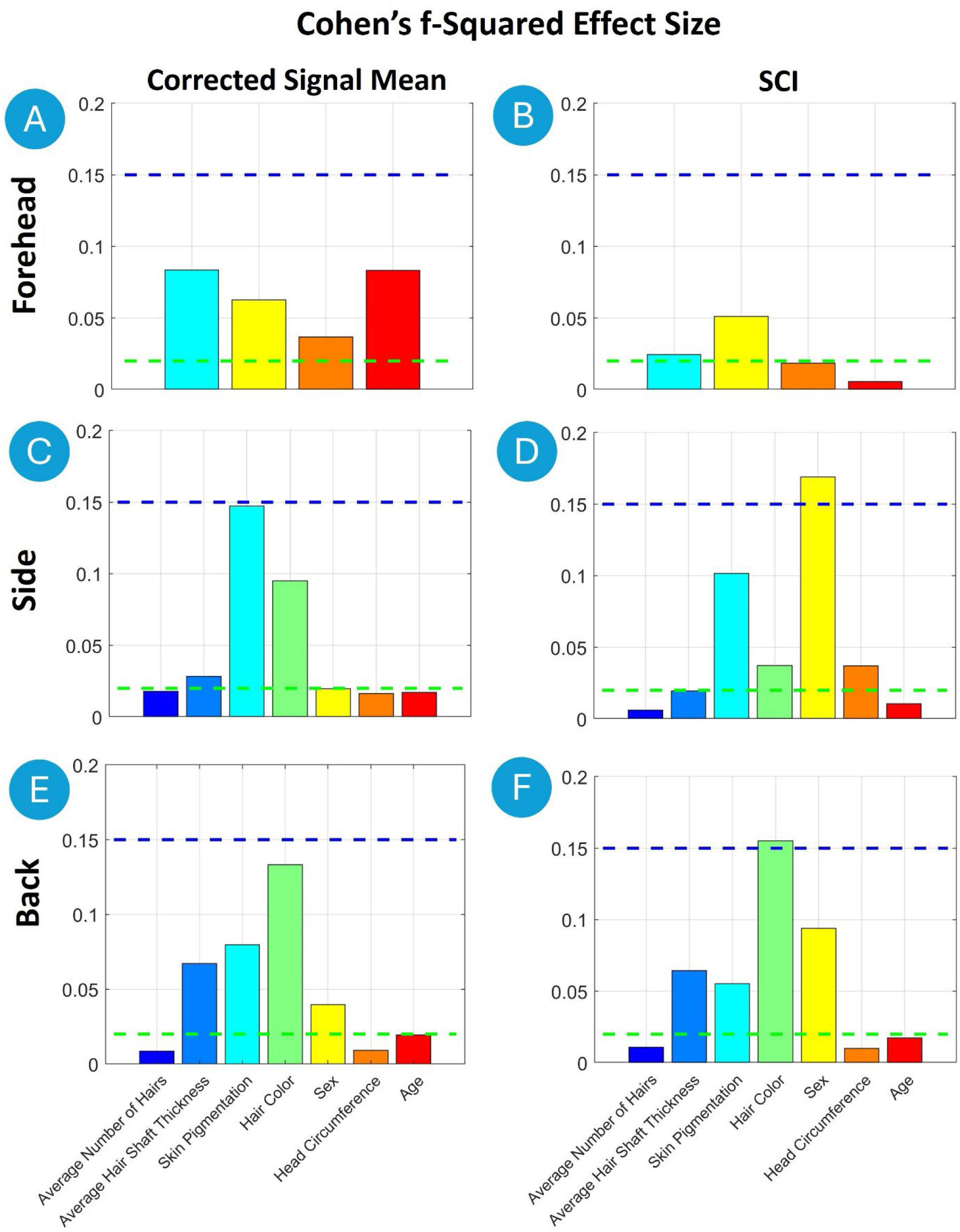
Cohen’s f² effect sizes from the multiple linear regression analysis predicting fNIRS-signal quality on the Forehead, Side of the Head and Back of the Head. Effect sizes were interpreted according to Cohen’s (1988) guidelines, where f² ≥ 0.02 (green horizontal line) indicates a small effect, f² ≥ 0.15 a medium effect (blue horizontal line), and f² ≥ 0.35 a large effect.

**Figure 9.**
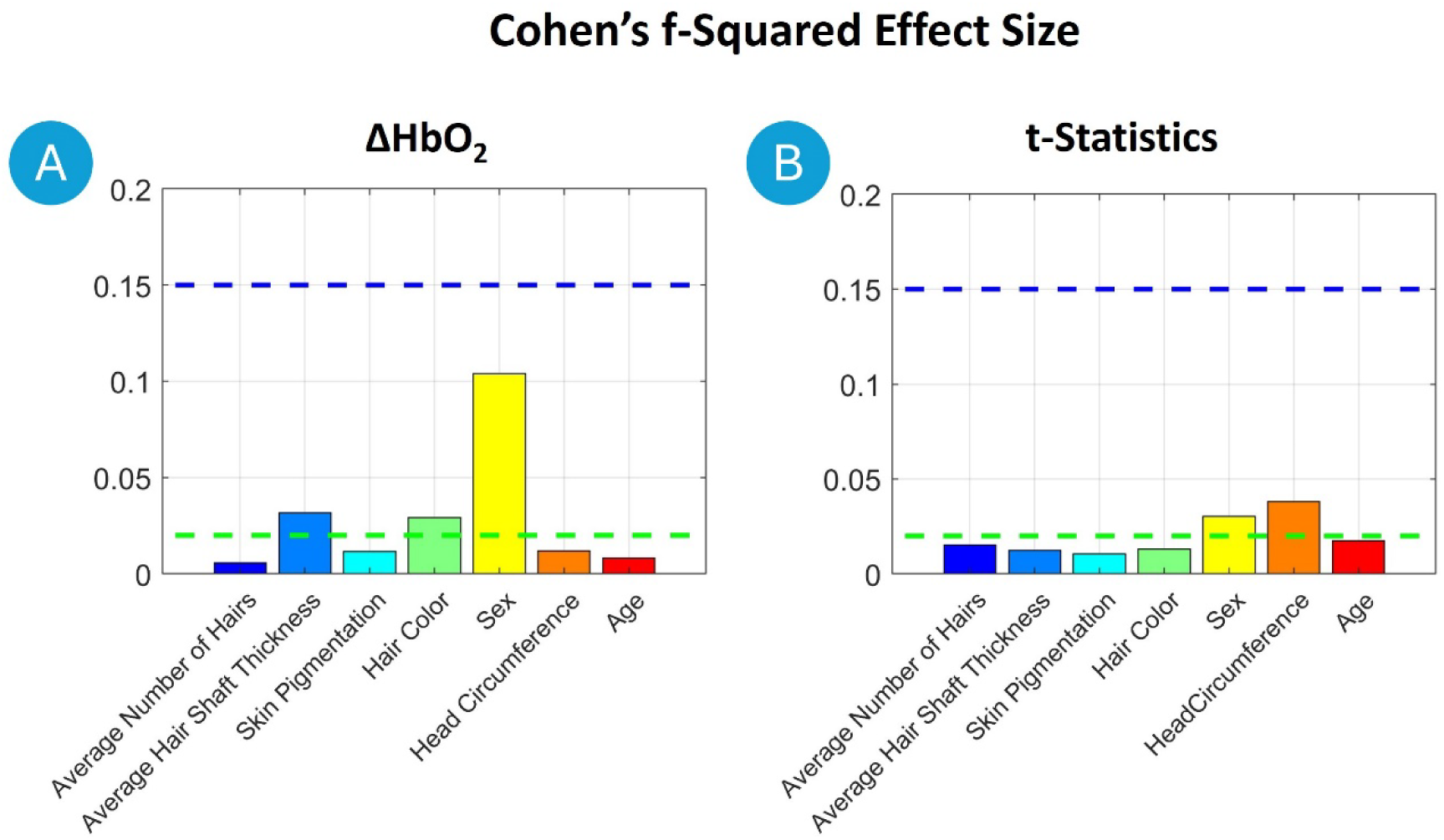
Cohen’s f² effect sizes from the multiple linear regression analysis predicting GLM output ΔHbO_2_ and t-statistics for a ball-squeezing task. Effect sizes were interpreted according to Cohen’s (1988) guidelines, where f² ≥ 0.02 (green horizontal line) indicates a small effect, f² ≥ 0.15 a medium effect (blue horizontal line), and f² ≥ 0.35 a large effect.

**Table 5:** Output of Multiple Linear Regression analysis for predictors of **Corrected Signal Mean** on the Forehead, Side of the Head and Back of the Head. The table presents the estimated coefficients, their standard errors (SE), t-statistics, p-values, -f-squared effect sizes (f^2^), and variance inflation factors (VIF) for each predictor variable in the model. Red highlights statistical significance of p <=0.01.

| Forehead | Estimate | SE | tStat | pValue | $f^2$ | VIF |
| --- | --- | --- | --- | --- | --- | --- |
| Skin Pigmentation | -0.009 | 0.002 | -6.2 | <0.01 | 0.08 | 1.1 |
| Sex | 0.384 | 0.058 | 6.7 | <0.01 | 0.06 | 1.2 |
| Head Circumference | 0.023 | 0.012 | 1.9 | 0.06 | 0.04 | 1.2 |
| Age | 0.015 | 0.003 | 6.0 | <0.01 | 0.08 | 1.0 |
| Side of the Head | Estimate | SE | tStat | pValue | $f^2$ | VIF |
| Average Number of Hairs | -0.001 | 0.000 | -2.6 | 0.01 | 0.02 | 1.2 |
| Average Hair Shaft Thickness | -0.016 | 0.003 | -6.0 | <0.01 | 0.03 | 1.2 |
| Skin Pigmentation | -0.020 | 0.001 | -17.5 | <0.01 | 0.15 | 1.2 |
| Hair Color | -0.290 | 0.022 | -13.3 | <0.01 | 0.10 | 1.4 |
| Sex | -0.049 | 0.042 | -1.2 | 0.25 | 0.02 | 1.3 |
| Head Circumference | -0.007 | 0.009 | -0.7 | 0.48 | 0.02 | 1.3 |
| Age | 0.006 | 0.002 | 2.9 | <0.01 | 0.02 | 1.4 |

|  | Estimate | SE | tStat | pValue | f<sup>2</sup> | VIF |
| --- | --- | --- | --- | --- | --- | --- |
| **Back of the Head** |  |  |  |  |  |  |
| Average Number of Hairs | -0.001 | 0.001 | -1.2 | 0.23 | 0.01 | 1.4 |
| Average Hair Shaft Thickness | -0.044 | 0.005 | -8.7 | <0.01 | 0.07 | 1.3 |
| Skin Pigmentation | -0.017 | 0.002 | -7.9 | <0.01 | 0.08 | 1.3 |
| Hair Color | -0.414 | 0.040 | -10.3 | <0.01 | 0.13 | 1.4 |
| Sex | -0.319 | 0.077 | -4.1 | <0.01 | 0.04 | 1.3 |
| Head Circumference | 0.029 | 0.017 | 1.7 | 0.09 | 0.01 | 1.3 |
| Age | 0.014 | 0.004 | 3.6 | <0.01 | 0.02 | 1.4 |

On the Forehead, Skin Pigmentation showed a significant negative effect on Corrected Signal Mean (t = −6.2, p < 0.01, f² = 0.08), indicating that higher pigmentation correlates with reduced signal strength. Additionally, Sex had a positive effect (t = 6.7, p < 0.01, f² = 0.06), with females exhibiting higher Corrected Signal Mean compared to males.

On the Side of the Head, Skin Pigmentation displayed the strongest negative effect on Corrected Signal Mean (t = −17.5, p < 0.01, f² = 0.15), followed by Hair Color (t = −13.3, p < 0.01, f² = 0.10). Darker pigmentation and hair color were associated with reduced signal quality. Average Hair Shaft Thickness also negatively affected Corrected Signal Mean (t = −6.0, p < 0.01, f² = 0.03). The Average Number of Hairs showed a weaker, but still significant, effect (t = −2.6, p = 0.01, f² = 0.02). Age had a positive effect (t = 2.9, p < 0.01, f² = 0.02), indicating that signal strength improved slightly with increasing age.

On the Back of the Head, Hair Color had the largest negative effect on the Corrected Signal Mean (t = −10.3, p < 0.01, f² = 0.13), followed by Skin Pigmentation (t = −7.9, p < 0.01, f² = 0.08). Average Hair Shaft Thickness also significantly reduced Corrected Signal Mean (t = −8.7, p < 0.01, f² = 0.07). Additionally, females had lower Corrected Signal Mean on the Back of the Head compared to males (t = −4.1, p < 0.01, f² = 0.04). Age showed a positive effect (t = 3.6, p < 0.01, f² = 0.02), suggesting improved signal with age.

Table 6 and Figure 8 summarize the results from the multiple linear regression analysis performed to identify predictors of SCI across all three regions.

**Table 6:** Output of Multiple Linear Regression Analysis for Predictors of **SCI** at the Forehead, Side of the Head and Back of the Head. The table presents the estimated coefficients, their standard errors, t-statistics, p-values, effect sizes (f-squared), and variance inflation factors for each predictor variable in the model. Red highlights statistical significance of p <=0.01.

| **Forehead** | Estimate | SE | tStat | pValue | f<sup>2</sup> | VIF |
| --- | --- | --- | --- | --- | --- | --- |
| Skin Pigmentation | -0.006 | 0.003 | -2.4 | 0.02 | 0.02 | 1.1 |
| Sex | -0.421 | 0.097 | -4.3 | <0.01 | 0.05 | 1.2 |
| Head Circumference | 0.054 | 0.021 | 2.6 | 0.01 | 0.02 | 1.2 |
| Age | 0.000 | 0.004 | 0.1 | 0.95 | 0.01 | 1.0 |

| **Side of the Head** | Estimate | SE | tStat | pValue | f<sup>2</sup> | VIF |
| --- | --- | --- | --- | --- | --- | --- |
| Average Number of Hairs | -0.001 | 0.001 | -0.9 | 0.37 | 0.01 | 1.2 |
| Average Hair Shaft Thickness | -0.015 | 0.004 | -3.9 | <0.01 | 0.02 | 1.2 |
| Skin Pigmentation | -0.018 | 0.002 | -10.8 | <0.01 | 0.10 | 1.2 |
| Hair Color | -0.185 | 0.032 | -5.7 | <0.01 | 0.04 | 1.4 |
| Sex | -0.854 | 0.062 | -13.7 | <0.01 | 0.17 | 1.3 |
| Head Circumference | -0.090 | 0.014 | -6.5 | <0.01 | 0.04 | 1.3 |
| Age | 0.007 | 0.003 | 2.3 | 0.02 | 0.01 | 1.4 |

Back of the Head
|  | Estimate | SE | tStat | pValue | f<sup>2</sup> | VIF |
| --- | --- | --- | --- | --- | --- | --- |
| Average Number of Hairs | -0.001 | 0.001 | -0.8 | 0.40 | 0.01 | 1.4 |
| Average Hair Shaft Thickness | -0.039 | 0.007 | -5.3 | <0.01 | 0.06 | 1.3 |
| Skin Pigmentation | -0.017 | 0.003 | -5.3 | <0.01 | 0.06 | 1.3 |
| Hair Color | -0.479 | 0.060 | -8.0 | <0.01 | 0.16 | 1.4 |
| Sex | -0.697 | 0.115 | -6.1 | <0.01 | 0.09 | 1.3 |
| Head Circumference | 0.016 | 0.026 | 0.6 | 0.52 | 0.01 | 1.3 |
| Age | 0.010 | 0.006 | 1.7 | 0.09 | 0.02 | 1.4 |

**Table 7:** Multiple linear regression analysis results for factors influencing GLM output changes in ΔHbO_2_ and t-statistics during the ball-squeezing task. The table displays the estimated coefficients, standard errors (SE), t-statistics of the regression, p-values, effect sizes f^2^, and variance inflation factors (VIF) for each predictor. Red highlights statistical significance of p <=0.01.

| $\Delta\text{HbO}_2$ | Estimate | SE | tStat | pValue | f_squared | VIF |
| --- | --- | --- | --- | --- | --- | --- |
| Average Number of Hairs | 0.000 | 0.000 | 1.0 | 0.33 | 0.01 | 1.2 |
| Average Hair Shaft Thickness | 0.000 | 0.000 | -6.3 | 0.00 | 0.03 | 1.2 |
| Skin Pigmentation | 0.000 | 0.000 | -1.7 | 0.09 | 0.01 | 1.2 |
| Hair Color | 0.000 | 0.000 | 5.4 | 0.00 | 0.03 | 1.4 |
| Sex | 0.000 | 0.000 | -10.6 | 0.00 | 0.10 | 1.3 |
| Head Circumference | 0.000 | 0.000 | -5.7 | 0.00 | 0.01 | 1.3 |
| Age | 0.000 | 0.000 | -3.3 | 0.00 | 0.01 | 1.4 |

| t-Statistics | Estimate | SE | tStat | pValue | f_squared | VIF |
| --- | --- | --- | --- | --- | --- | --- |
| Average Number of Hairs | 0.039 | 0.007 | 5.7 | <0.01 | 0.02 | 1.2 |
| Average Hair Shaft Thickness | -0.067 | 0.036 | -1.9 | 0.06 | 0.01 | 1.2 |
| Skin Pigmentation | -0.007 | 0.016 | -0.4 | 0.67 | 0.01 | 1.2 |
| Hair Color | 1.154 | 0.299 | 3.9 | <0.01 | 0.01 | 1.4 |
| Sex | -4.021 | 0.578 | -7.0 | <0.01 | 0.03 | 1.3 |
| Head Circumference | -0.812 | 0.129 | -6.3 | <0.01 | 0.04 | 1.3 |
| Age | 0.098 | 0.029 | 3.3 | <0.01 | 0.02 | 1.4 |

On the Forehead, the strongest predictor was Sex (t = −4.3, p < 0.01, f² = 0.05), with males showing higher SCI values compared to females. Head Circumference also had a positive effect (t = 2.6, p = 0.01, f² = 0.02), suggesting improved coupling with larger head sizes on the Forehead.

On the Side of the Head, the largest effect was from Sex (t = −13.7, p < 0.01, f² = 0.17), with males exhibiting higher SCI values than females. Skin Pigmentation also had a substantial negative effect (t = −10.8, p < 0.01, f² = 0.10), followed by Hair Color (t = −5.7, p < 0.01, f² = 0.04). The Head Circumference was also negatively associated with SCI (t = −6.5, p < 0.01, f² = 0.04) as well as the Average Hair Shaft (t = −3.9, p < 0.01, f² = 0.02).

On the Back of the Head, Hair Color had the largest effect (t = −8.0, p < 0.01, f² = 0.16), followed by Sex (t = −6.1, p < 0.01, f² = 0.09), males having larger SCI values than female, and Skin Pigmentation (t = −5.3, p < 0.01, f² = 0.06). Increased Average Hair Shaft Thickness also reduced SCI (t = −5.3, p < 0.01, f² = 0.06).

### 3.5 Ball-Squeezing Task

In terms of ΔHbO_2_, Sex emerged as the most significant predictor, male having higher ΔHbO_2_ than female (t = −10.6, p < 0.01, f² = 0.10). Average Hair Shaft Thickness had a significant negative impact (t = −6.3, p < 0.01, f² = 0.03), suggesting that thicker hair is associated with lower ΔHbO_2_ levels. Additional predictors included Head Circumference, which negatively influenced ΔHbO_2_ (t = −5.7, p < 0.01, f² = 0.01), and Age, which also showed a notable negative effect (t = −3.3, p < 0.01, f² = 0.01). Hair Color had a significant positive effect on ΔHbO_2_ (t = 5.4, p < 0.01, f² = 0.03).

For the t-statistics, Sex was again the most influential factor, male having higher t-statistics than female (t = −7.0, p < 0.01, f² = 0.03). The Head Circumference significantly affected t-statistics as well (t = −6.3, p < 0.01, f² = 0.04), implying reduced t-values with increased head circumference. Average Number of Hairs had a positive effect (t = 5.7, p < 0.01, f² = 0.02) on t-statistics. Finally, Age positively influenced t-statistics (t = 3.3, p < 0.01, f² = 0.02).

### 3.6 Effect of Increased Optode-Scalp Coupling through Intensive Hair Adjustment

Our study included two resting-state runs in order to see the improvement obtained with thorough hair adjustment. The first run was started with only minimal hair adjustment (< 1 min; i.e., fast capping), and the second run was started after a thorough hair adjustment (i.e., proper capping). Table 8 summarizes the improvement with additional hair adjustment.

**Table 8:** Mean, standard deviation and the percent change from run 1 to Run 2 of Uncorrected Signal Mean and SCI for the Forehead, Side of the Head and Back of the Head.

| ROI | Measure | Run 1<br>Mean (SD) | Run 2<br>Mean (SD) | Percent<br>Increase (%) | P-Value |
| --- | --- | --- | --- | --- | --- |
| Forehead | SCI | 0.89 (0.21) | 0.95 (0.10) | 6.0 | <0.01 |
|  | Uncorrected Signal Mean | 0.33 (0.10) | 0.33 (0.10) | 0.7 | 0.79 |
| Side | SCI | 0.78 (0.27) | 0.90 (0.18) | 15.4 | <0.01 |
|  | Uncorrected Signal Mean | 0.14 (0.12) | 0.17 (0.11) | 18.8 | <0.01 |
| Back | SCI | 0.63 (0.35) | 0.76 (0.29) | 20.7 | <0.01 |
|  | Uncorrected Signal Mean | 0.10 (0.15) | 0.11 (0.13) | 9.2 | 0.21 |

For the Forehead, the mean SCI increased from Mean (SD) of 0.89 (0.21) in Run 1 to 0.95 (0.10) in Run 2, showing a significant percent increase of 6% (Table 8). In contrast, there was no statistically significant change in Uncorrected Signal Mean.

On the Side of the Head, the SCI measure showed a significant improvement, with the mean increasing from 0.78 (0.27) in Run 1 to 0.90 (0.19) in Run 2, representing a 15.4% increase. Similarly, the Uncorrected Signal Mean also demonstrated a significant increase, with the mean rising from 0.14 (0.19) in Run 1 to 0.17 (0.11) in Run 2, corresponding to an 18.8% increase.

On the Back of the Head, the SCI measure exhibited a significant improvement, with the mean value increasing from 0.63 (0.36) in Run 1 to 0.76 (0.29) in Run 2, i.e. 20.7% increase. In contrast, there was no statistically significant improvement in the Uncorrected Signal Mean.

### 3.7 Inclusivity with Noise Equivalent Power (NEP) levels

Figure 10A shows the distribution of Uncorrected Signal Mean in each channel against a combined hair-skin metric across all participants in the study. For the sake of further analysis, we combined hair and skin characteristics (hair color, type, and thickness and skin pigmentation) into a single parameter which we call a combined hair-skin metric. A higher value of this metric indicates more challenging fNIRS acquisition due to higher light absorption by hair and skin. Binning data points yields five groups that range from a very low (0.15-0.30) to a very high (0.75-1) combined hair-skin metric. Of all channels, these groups represent 9.7% (Group 1, [0.15-0.30]), 31.8% (Group 2, [0.30-0.45]), 41.1% (Group 3, [0.45-0.60]), 9.3% (Group 4, [0.60.0.75]) and 8.1% (Group 4, [0.75-1]) of the participants measured. The overall distribution shows a clear trend towards lower light levels for higher combined hair-skin metric skin.

The amount of measured light in the presence of a higher combined hair-skin metric is influenced by the sensitivity of the detection branch of an fNIRS device. Consequently, the sensitivity of the detectors— a fundamental performance characteristic of the device—can be related to the measured and expected number of optimal channels. Utilizing our technical knowledge of the sensitivity of the NIRSport2 device employed in this study, we relate the measured raw signal noise floor of the device to its equivalent detector sensitivity, known as Noise Equivalent Power (NEP) (here: 6.8^−6^ ≍ 52 *fW* ∕ √*Hz*). This equivalence allows us to estimate the performance of fNIRS systems with a Noise Equivalent Power (NEP) that is equal to or greater than that of the measurement system. By virtually raising the system’s NEP threshold and establishing a desired signal-to-noise ratio (SNR) of at least 20 dB—below which channels would typically be pruned—we can estimate the percentage of suboptimal channels in each group based on system sensitivity (NEP).

In our study with the given system, the NEP+20dB threshold yields a total of 5% suboptimal channels across all subjects. The relationship between the number of channels within each group relative to the system sensitivity is further detailed in Figure 10B, which highlights the importance of good detector sensitivity for inclusivity in fNIRS studies. A few examples are given as follows using again a 20dB threshold for SNR-based channel pruning. For a system with 50fW/sqrt(Hz) detector sensitivity, 100% of all channels in Group 1 pass the quality criterion, 96% of Group 2, 97% of Group 3, 91% of Group 4 and 89% of Group 5. In contrast, for fNIRS systems with a higher NEP of 1pW/sqrt(Hz), 98% of all channels pass for Group 1, 88% of Group 2, 74% of Group 3, down to 65% of Group 4, and only 50% of Group 5.

**Figure 10.**
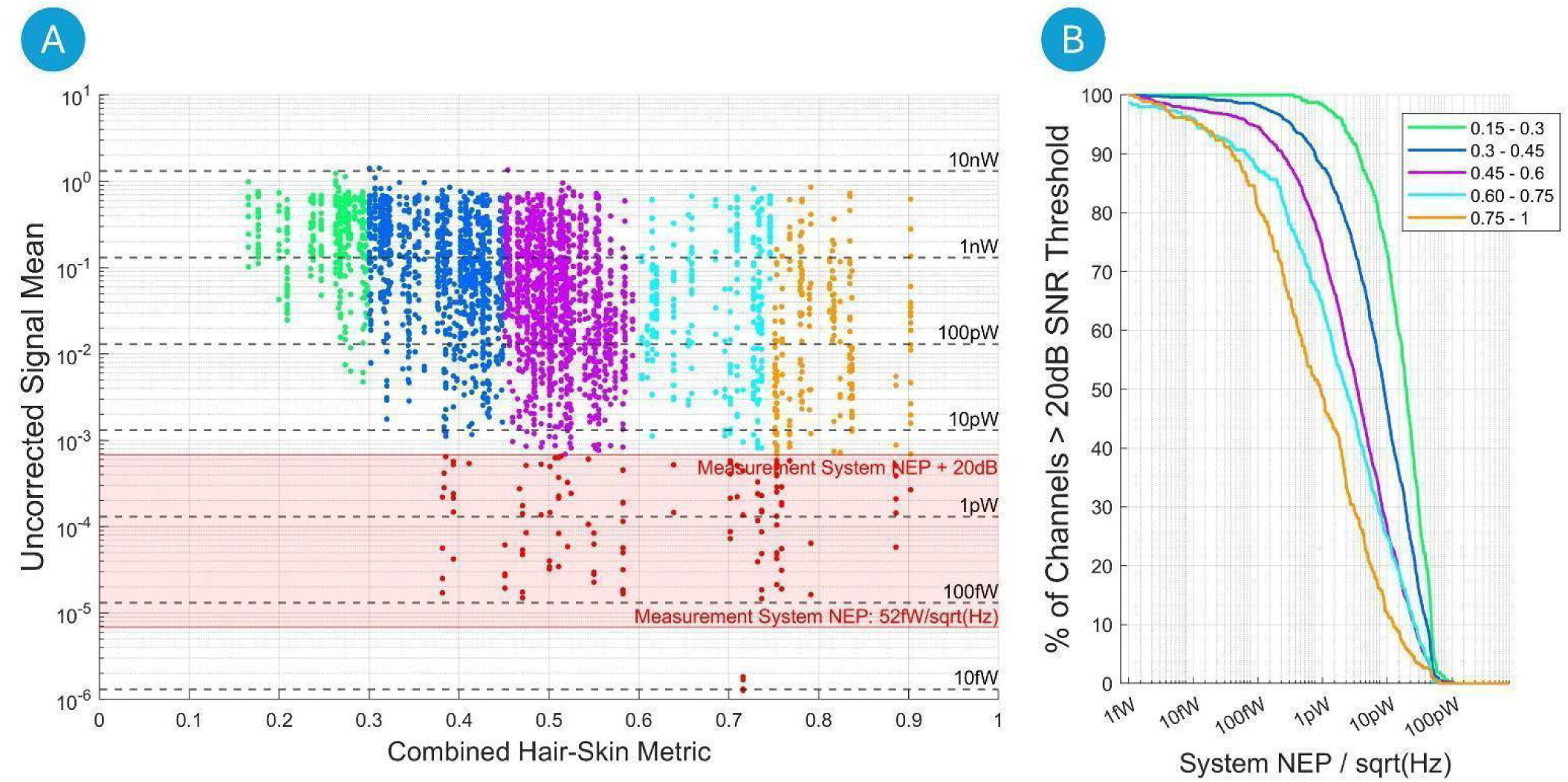
**A)** Scatter plot of Uncorrected Signal Mean in individual channels across all participants versus the Combined Hair-Skin Metric. The y-axis is in logarithmic scale. Grey dashed horizontal lines represent noise equivalent power levels (NEP/√(Hz)), beneath which any signal drowns in measurement noise. The measurement system’s NEP is 52fW/√(Hz) at an equivalent signal input noise level of 6.8*10^-6^ and a detector area of 9.6 mm². The red area highlights all channels that must be omitted if an SNR signal quality threshold of two orders of magnitude (20dB) above the NEP is applied. Example: In a system with an NEP of 1pW, with a 20dB threshold, no channel below the vertical line at 100pW would pass the quality constraint. This is further quantified in subplot **B)** For each binned group of Combined Hair-Skin Metric, the plot shows the percentage of channels that pass a 20dB signal quality threshold relative to system sensitivity on the x-axis. Example: For the third group in purple color (i.e. Combined Hair-Skin Metric ∈ [0.45-0.6]), 98% of all channels would pass the quality threshold for a system with sensitivity of 10fW/sqrt(Hz), but only 70% of all channels would pass the threshold for a system with sensitivity of 1pW/sqrt(Hz).

## 4 Discussion

The present study explored various predictors of fNIRS signal quality across three head regions: Forehead, Side of the Head, Back of the Head. A key finding from our analysis was the consistent modulatory effect of hair properties on signal intensity, particularly on the Side and Back of the Head. Hair color, hair type (e.g., straight, curly, or kinky), hair strand texture (e.g., fine, medium, coarse), and hair shaft thickness are critical factors that can significantly reduce the quality of optical signals in fNIRS measurements. These characteristics may either increase light absorption or obstruct the transmission of signals. For instance, darker hair colors typically absorb more light, thereby diminishing the amount of light that reaches the detectors, and thicker and coarser hair can physically block and/or scatter light. Significant advancements in the field of fNIRS are necessary, which may include innovative strategies or modifications of established techniques, such as incorporating brush-fiber tips to enhance optode-scalp coupling ^29^. In addition to optimizing optode design, exploring flexible and adaptable methods for hair management, including custom-fit caps ^18^ or alternative optical interfaces, may enhance signal acquisition for individuals with various hair characteristics.

Hair style, which refers to how hair is arranged or modified (e.g., braids, buns, twists, locs, or ponytails), is another potential factor that can affect optode-scalp coupling, and thus signal quality. Due to the numerous hair styles and the resulting categorical variables, we chose not to include this factor in our analysis, as our limited sample size would not adequately support modeling it. However, it is important to note that there are ongoing efforts in both the EEG and fNIRS fields to address this issue. For example, techniques such as parted braiding allow for better alignment of electrodes or optodes with hair parts, reducing overall volume while maintaining hair-free sections ^35–37^.

Skin pigmentation was also identified as a significant factor. Our results showed that higher skin pigmentation is associated with reduced signal intensity as well as SCI. This phenomenon can be attributed to the increased absorption of the signal by darker pigments. Parallels can be drawn with the pulse oximetry field, where there has been a growing societal and governmental push to enhance the inclusivity and accuracy of devices that utilize near-infrared light for clinical measurements. Pulse oximetry is a widely used technique for measuring absolute levels of oxygen saturation in arterial blood^38^ and relies on the differential absorption of red and infrared light by oxygenated and deoxygenated hemoglobin. Although highly effective, pulse oximetry’s accuracy can be compromised by skin pigmentation due to melanin’s absorption characteristics. Higher melanin levels result in greater light absorption, which can interfere with the accurate detection of transmitted or reflected light, particularly at lower oxygen saturation levels. Studies, such as those by Feiner et al. (2007), have shown that darker skin can lead to significant inaccuracies in pulse oximeter readings ^39^, especially at low oxygen saturation levels, often resulting in undetected hypoxemia ^40,41^. However, it is important to note that pulse oximetry measures absolute levels of oxygen saturation in arterial blood. In contrast, continuous-wave functional near-infrared spectroscopy (CW fNIRS) measures relative changes in oxygenated and deoxygenated hemoglobin concentrations, rather than providing absolute values, and generally measures deeper tissues such that skin pigmentation is a much smaller contributor to signal attenuation and thus the issue for CW fNIRS is a reduced signal quality rather than accuracy.

On a related note, skin type, as classified by the Fitzpatrick Skin Phototype Scale ^23^, was found to be significantly correlated with skin pigmentation measurements; therefore, it was not included in our regression models. We recommend using a Melanometer for an objective assessment of skin pigmentation. However, for researchers who do not have access to a Melanometer, we suggest documenting skin type as assessed by the Fitzpatrick Skin Phototype Scale. This approach allows for a standardized reference point, ensuring that the variability in skin type is acknowledged even in the absence of objective measurements.

Sex also emerged as a significant predictor across various metrics. On the forehead, females exhibit higher Corrected Signal Mean, likely due to thinner skulls ^42^ and less dense bone structure ^43^, which reduce signal attenuation and enhance optical penetration. However, at the side and back of the head, females show lower Corrected Signal Mean, possibly due to hair length, and styles (e.g., ponytails, braids) that were not accounted for in the model but could affect light transmission. The reduction in SCI from males to females across all regions may be attributed to physiological differences, such as males having larger heart sizes, greater aerobic capacity, and higher cardiac output^44^, contributing to the observed SCI differences. However, the difference between males and females is much less pronounced in the t-statistics. This is likely because t-statistics account for both the magnitude of the signal (contrast) and its variability (noise), offering a measure of contrast-to-noise ratio rather than just the contrast. As such, even if males generate higher ΔHbO_2_ signals, intra-subject variability could reduce the statistical distinction in the t-statistics.

For the ΔHbO_2_ contrast and t-statistics, males displayed larger values than females during the ball-squeezing task. This may reflect the influence of greater muscle mass and strength in male ^45^. As a result, male participants might have engaged more muscle fibers during the task, leading to increased blood flow and oxygen delivery to the active tissues, and thereby generating higher ΔHbO_2_. The difference is less drastic in t-statistics, highlighting the importance of statistical analysis that accounts for the variance.

As individuals age, they typically experience changes in hair density and color, which can positively affect signal quality. While age was significantly correlated with the Corrected Signal Mean, it did not emerge as a significant predictor in our regression model, which accounted for hair characteristics, thus masking the age effect itself. Interestingly, age was a predictor of higher Corrected Signal Mean specifically in the forehead region. This observation may be explained by physiological changes associated with aging.

Our results show that low Noise Equivalent Power (i.e., high system sensitivity) is crucial for a balanced distribution of channels from subjects with varying skin and hair characteristics. For more inclusive fNIRS studies, a significant number of channels must pass signal quality. While our analysis highlights that a high number of optimal channels is not always a given and strongly depends on system performance, they also show that state-of-the-art equipment with highly sensitive detectors can indeed provide the required performance. Further advancements to reduce the impact of hair-related variables, such as improved sensor placement and innovative optode designs could enhance data quality even more and allow more inclusive research. For instance, advancements in sensor technology that allow better penetration through dense hair ^46^ or adaptive algorithms that adjust to varying hair conditions could improve signal acquisition, leading to more reliable and interpretable results.

We provide a metadata recommendation table (Table 9) for fNIRS researchers, emphasizing the importance of documenting participant-level factors that influence the quality and interpretability of fNIRS signals. Including this metadata in Brain Imaging Data Structure (BIDS) format ^47,48^ within open repositories like OpenNeuro enhances inclusivity by ensuring that diverse populations are accurately represented in neuroimaging research. Systematic documentation of these factors using standardized scales—such as the Fischer-Saller scale for hair color, the Andre Walker system for hair type, the FIA for hair strand texture, and the Fitzpatrick scale for skin type—will enable researchers to identify and address potential sources of variability in optical signal attenuation. Collecting demographic data such as age, sex, and ethnicity enriches the dataset and allows for a more comprehensive analysis of how these factors interact to affect signal quality. Optional yet beneficial metadata, including hair measurements via a Trichoscope and skin pigmentation (Melanin Index) measurements with a Melanometer, can provide valuable insights into individual variability. Ultimately, the inclusion of these metadata in more openly distributed BIDS compliant datasets will enable meta-analyses of the impact of these participant-level factors on fNIRS recordings from large populations of subjects.

**Table 9:**
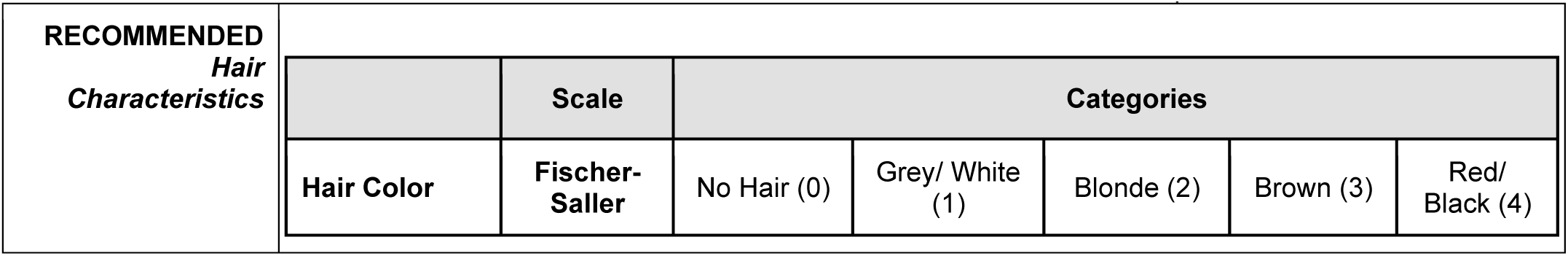
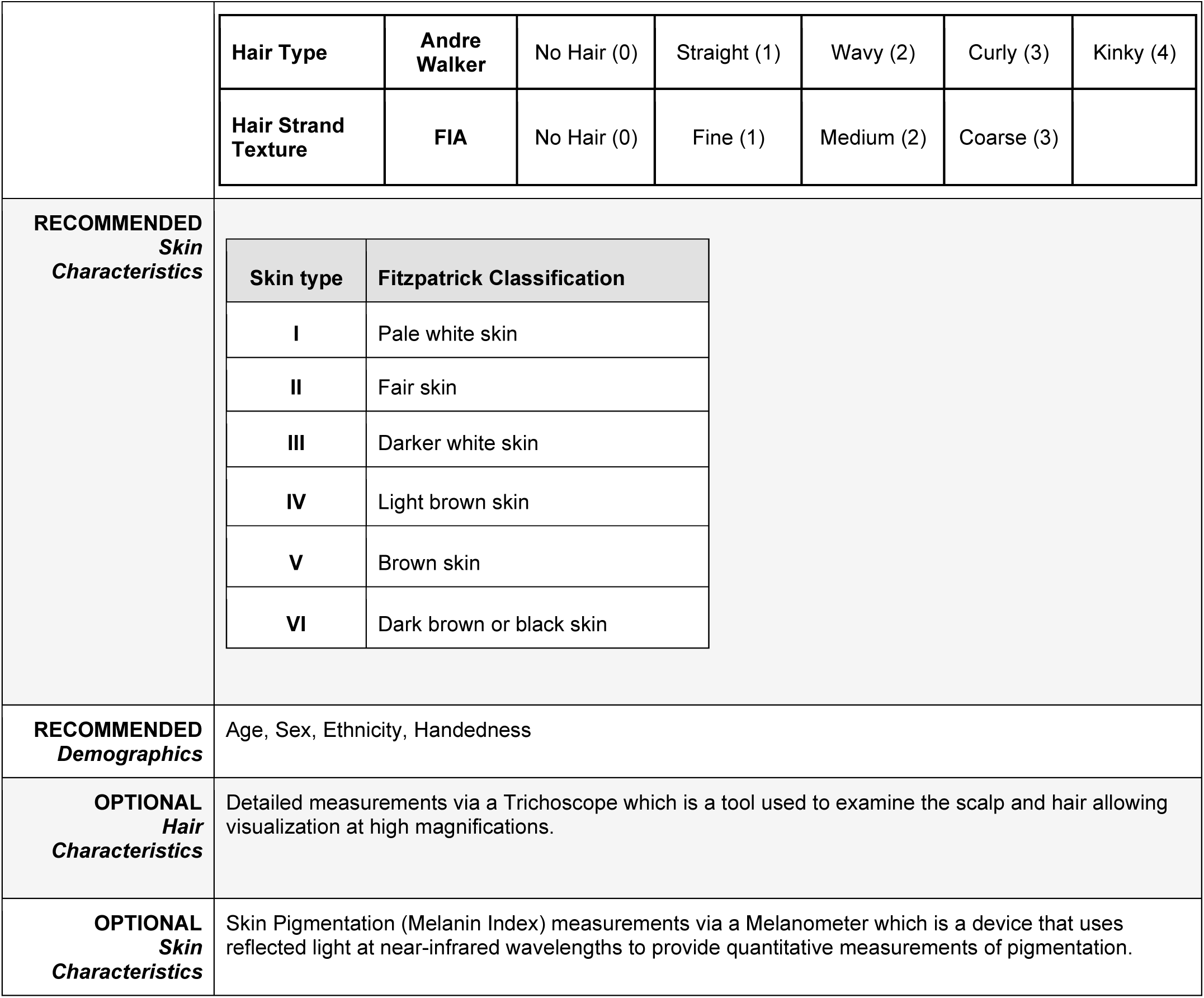
Metadata collection recommendations for fNIRS researchers with recommended and optional fields.

Table 10 presents a comprehensive overview of best practices and recommendations for optimizing fNIRS data collection, taking into account diverse participant-level factors. It underscores the importance of effective communication with participants, focusing on collaborative strategies to enhance signal quality without placing blame on the individual. The table presents key advancements in cap design, such as the innovative 3D-printed NinjaCaps ^18^, alongside future directions, including the development of wireless designs that enhance participant mobility by eliminating cumbersome wires. The table also highlights specific hair adjustment techniques and specialized optode placement strategies to maximize scalp exposure. Furthermore, it discusses the role of chin straps and real-time feedback in ensuring cap stability and consistent signal quality, alongside hygienic considerations for fieldwork using easily disinfectable materials.

**Table 10:**
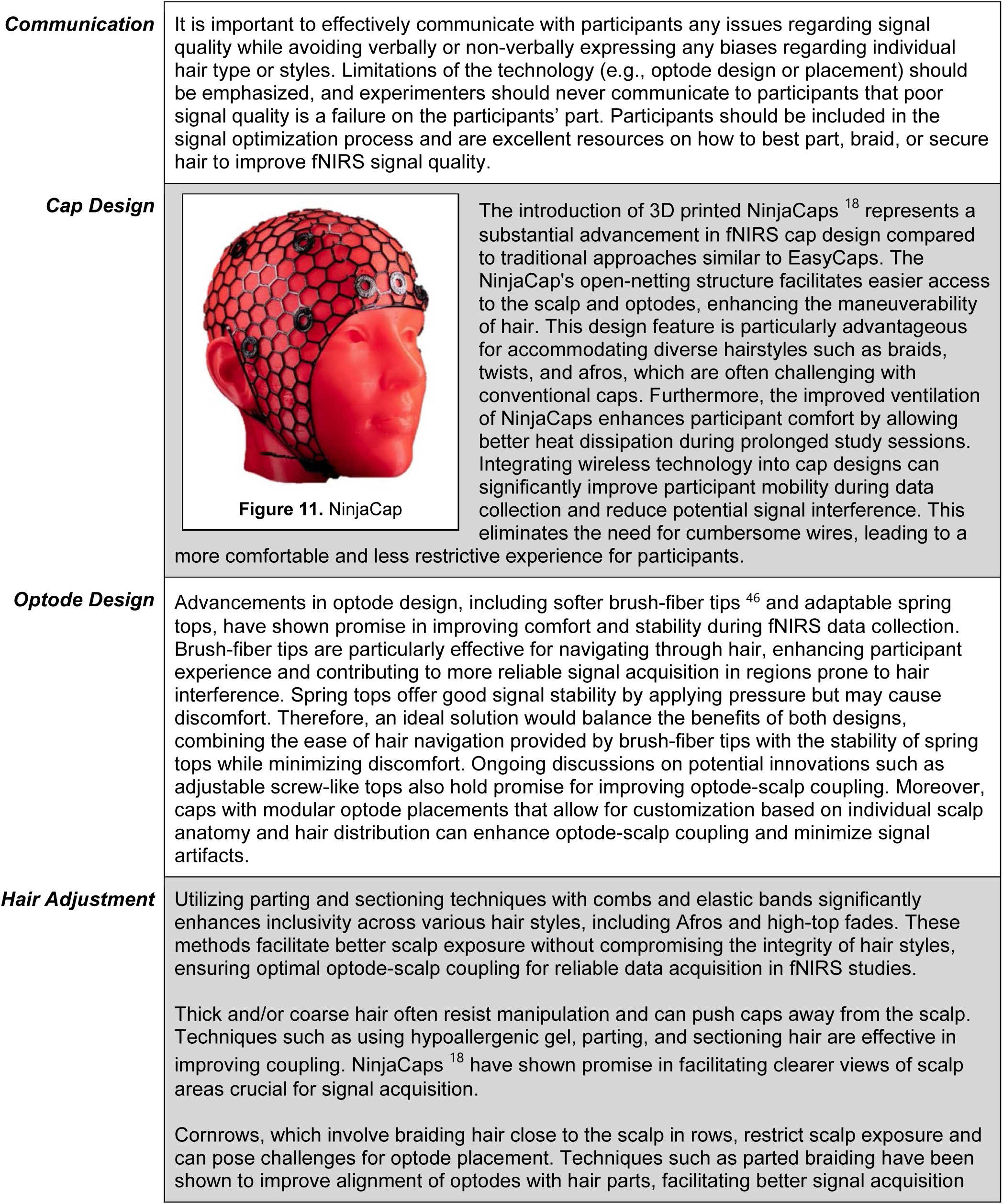
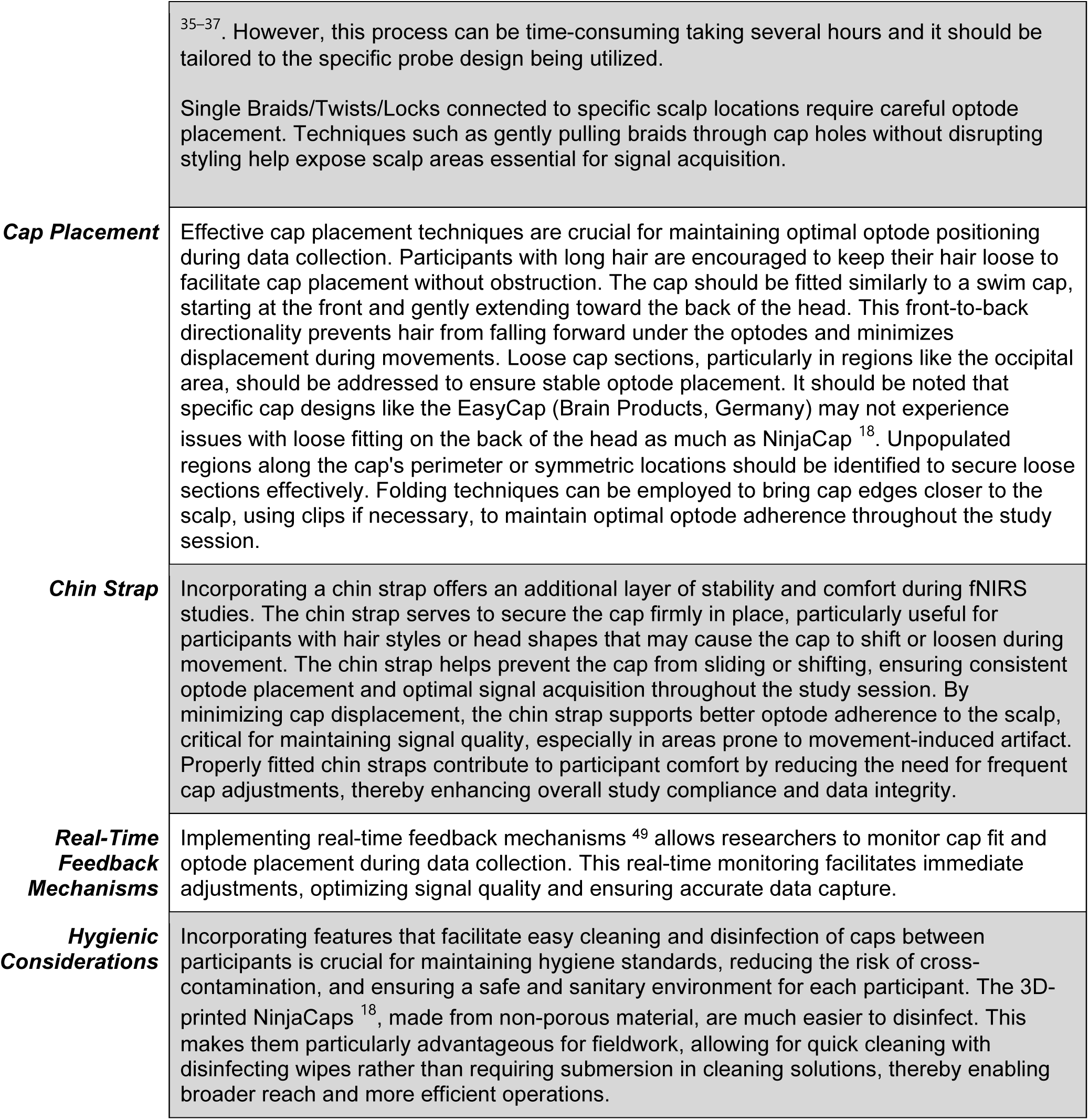
Best practices and recommendations for optimizing fNIRS data collection across diverse participant characteristics.

### Communication Cap Design Optode Design Hair Adjustment

It is important to effectively communicate with participants any issues regarding signal quality while avoiding verbally or non-verbally expressing any biases regarding individual hair type or styles. Limitations of the technology (e.g., optode design or placement) should be emphasized, and experimenters should never communicate to participants that poor signal quality is a failure on the participants’ part. Participants should be included in the signal optimization process and are excellent resources on how to best part, braid, or secure hair to improve fNIRS signal quality.

The introduction of 3D printed NinjaCaps ^18^ represents a substantial advancement in fNIRS cap design compared to traditional approaches similar to EasyCaps. The NinjaCap’s open-netting structure facilitates easier access to the scalp and optodes, enhancing the maneuverability of hair. This design feature is particularly advantageous for accommodating diverse hairstyles such as braids, twists, and afros, which are often challenging with conventional caps. Furthermore, the improved ventilation of NinjaCaps enhances participant comfort by allowing better heat dissipation during prolonged study sessions. Integrating wireless technology into cap designs can significantly improve participant mobility during data collection and reduce potential signal interference. This eliminates the need for cumbersome wires, leading to a more comfortable and less restrictive experience for participants.

Advancements in optode design, including softer brush-fiber tips ^46^ and adaptable spring tops, have shown promise in improving comfort and stability during fNIRS data collection. Brush-fiber tips are particularly effective for navigating through hair, enhancing participant experience and contributing to more reliable signal acquisition in regions prone to hair interference. Spring tops offer good signal stability by applying pressure but may cause discomfort. Therefore, an ideal solution would balance the benefits of both designs, combining the ease of hair navigation provided by brush-fiber tips with the stability of spring tops while minimizing discomfort. Ongoing discussions on potential innovations such as adjustable screw-like tops also hold promise for improving optode-scalp coupling. Moreover, caps with modular optode placements that allow for customization based on individual scalp anatomy and hair distribution can enhance optode-scalp coupling and minimize signal artifacts.

Utilizing parting and sectioning techniques with combs and elastic bands significantly enhances inclusivity across various hair styles, including Afros and high-top fades. These methods facilitate better scalp exposure without compromising the integrity of hair styles, ensuring optimal optode-scalp coupling for reliable data acquisition in fNIRS studies.

Thick and/or coarse hair often resist manipulation and can push caps away from the scalp. Techniques such as using hypoallergenic gel, parting, and sectioning hair are effective in improving coupling. NinjaCaps ^18^ have shown promise in facilitating clearer views of scalp areas crucial for signal acquisition.

Cornrows, which involve braiding hair close to the scalp in rows, restrict scalp exposure and can pose challenges for optode placement. Techniques such as parted braiding have been shown to improve alignment of optodes with hair parts, facilitating better signal acquisition

Reporting participant-level factors in fNIRS studies is important, as these variables can influence signal acquisition and introduce variability that may affect results if not addressed. Excluding data based on specific hair and skin pigmentation can lead to biased outcomes, particularly when comparing groups with differing characteristics. For example, to maintain equal sample sizes for analyzable data, researchers should consider recruiting more participants for groups likely to experience unbalanced pruning. Our findings also indicate that sex can influence the contrasts obtained during functional tasks, suggesting that caution is warranted when comparing groups of different sexes. Integrating these variables into regression models as covariates may help control for their effects on the results.

## 5 Conclusion

Our study provides valuable insights into the participant-level predictors of signal quality in fNIRS neuroimaging. Hair characteristics, skin pigmentation, sex, and age emerged as significant factors affecting signal quality. By acknowledging and addressing these factors, researchers can enhance the inclusivity of neuroimaging technologies, paving the way for more equitable and effective applications in cognitive neuroscience and clinical diagnostics and treatment.

## 6 Data and Code Availability

The data used in this study is available at BfNIRS https://www.bfnirs.openfnirs.org/datasets, under the name “Inclusion Study”. The code used for data analysis and figure generation is available on GitHub at https://github.com/mayucel/InclusionStudy.

## 7 Acknowledgements

This research was supported by Meta Reality Labs (formerly Facebook Technologies, LLC) as part of the Engineering Approaches to Responsible Neural Interface Design Initiative (MAY). We would also like to acknowledge NIH U01EB0239856 (MAY, DB, SK, DS, ACG, TDE), NSF Research Traineeship Program (DGE-1633516) (EC), the Netherlands Organization for Scientific Research (NWiOd i-VGrant VI.Vidi.191.210) (BS), and the German Federal Ministry of Education and Research (BIFOLD24B) (AvL). We thank the NIRx team for their valuable support in guiding us on source power correction and helping us locate the relevant information in the acquired data.

## 8 Author Contributions

MAY and DAB conceptualized the research question and framework. MAY led the project. MAY, DAB, JEA, DR, and YG designed the experimental approach and protocols. PH, PF, and NM managed the recruitment of participants. JEA, DR, PH, PF, RIK, EJB, NM, SD, LC, DB, LKB, EC, JG, JW, VT, and YZ executed the experiments. MAY conducted data analysis and statistical assessments. MAY and AvL created the visualizations of the findings. MAY drafted the original manuscript. MAY, DAB, BS, AvL, EC, JEA, DJR, RIK, EJB, NM, ACG, and TDE reviewed, provided feedback and edited the manuscript. MAY secured funding for the research project. All authors approved the final version of the paper.

